# *In Silico* Electrophysiology of Inner-Ear Mechanotransduction Channel TMC1 Models

**DOI:** 10.1101/2021.09.17.460860

**Authors:** Sanket Walujkar, Jeffrey M. Lotthammer, Collin R. Nisler, Joseph C. Sudar, Angela Ballesteros, Marcos Sotomayor

## Abstract

Inner-ear sensory hair cells convert mechanical stimuli from sound and head movements into electrical signals during mechanotransduction. Identification of all molecular components of the inner-ear mechanotransduction apparatus is ongoing; however, there is strong evidence that TMC1 and TMC2 are pore-forming subunits of the complex. We present molecular dynamics simulations that probe ion conduction of TMC1 models built based on two different structures of related TMEM16 proteins. Unlike most channels, the TMC1 models do not show a central pore. Instead, simulations of these models in a membrane environment at various voltages reveal a peripheral permeation pathway that is exposed to lipids and that shows cation permeation at rates comparable to those measured in hair cells. Furthermore, our analyses suggest that TMC1 gating mechanisms involve protein conformational changes and tension-induced lipid-mediated pore widening. These results provide insights into ion conduction and activation mechanisms of hair-cell mechanotransduction channels essential for hearing and balance.

## INTRODUCTION

In mammalian hearing, pressure waves from sound are transmitted through the auditory canal to sensory hair cells responsible for mediating auditory perception in the cochlea of the inner ear^1–4^. Similarly, in the vestibular system, forces from head movements are conveyed to vestibular hair cells responsible for the sense of balance^5,6^. The hair-cell mechanosensory organelle consists of a bundle of hair-like projections called stereocilia that are arranged in rows of increasing height. Proteinaceous oblique tip links connect the tip of each stereocilium to the side of the taller stereocilium neighbor^7–9^ and are tensioned when mechanical stimuli cause deflections of hair bundles towards their tallest stereocilia row. This tension is then transferred to a mechanically activated transduction channel complex that facilitates an influx of potassium ions, depolarizing the hair cell^10^. This process occurs over microsecond timescales and triggers rapid signaling cascades responsible for auditory and balance perception^11,12^.

Determining the molecular identity of the inner-ear mechanotransduction channel has been difficult. There is an average of about 10,000 hair cells in the vertebrate inner ear, which is orders of magnitude smaller than the number of receptors in other sensory systems that have been better characterized^13^. Experiments also suggest that there are about two channels per tip link, with about 50 to 200 tip links per hair cell^14,15^. This scarcity of protein makes the complex difficult to study by traditional biochemical techniques^16^, explaining why the molecular identity of the mechanotransduction channel protein(s) has remained elusive for decades^13,17,18^. Nonetheless, exquisite electrophysiological and biophysical experiments have determined the force-dependent activation, localization, conductance, and selectivity of the hair-cell mechanotransduction channel in *ex vivo* tissue explants^10–12,15,19–23^. These data place at least four constraints on protein candidates for the channel. The protein(s) must be essential for hearing and force activated, be localized to the base of the tip link, and expressed at the appropriate stage during inner-ear development. Last, the mechanotransduction channel protein(s) must have a large cation selective pore with a single-channel conductance similar to that measured for the mechanotransduction channel *ex vivo* of approximately 50-180 pS^23,24^.

Several potential channel candidates have been proposed throughout the past two decades, including members of the TRP, HCN, ENaC, and ASIC families of proteins^25–40^. Despite the initial promise of each of these candidates, they all have been dismissed as none met all criteria listed above. However, genetic analyses and subsequent biochemical studies have shown that at least two members of the transmembrane channel-like (TMC) family of proteins, TMC1 and TMC2, as well as three additional membrane proteins, are part of the mechanotransduction complex^41–51^. These proteins include the lower part of the tip link made of PCDH15 with its transmembrane domain^47,52,53^, the tetraspan membrane protein of hair-cell stereocilia (TMHS, also known as LHFPL5)^49^, and the transmembrane inner-ear expressed protein (TMIE)^50,51^. All are essential for hearing, are localized at the base of the tip links, and might form a complex^43,46,48–52^. Yet, which of these, if any, forms the pore of the mechanotransduction complex remains controversial^13,17,18^.

The structure of TMHS in complex with PCDH15’s transmembrane domain and part of the PCDH15 ectodomain revealed a dimer of dimers that is unlikely to form an ion-conducting pore^54^, ruling out TMHS and PCDH15 as pore-forming subunits of the mechanotransduction complex. TMIE is predicted to have a single transmembrane helix^50,55,56^ and thus unlikely to form an ion-conducting pore by itself, although there is some evidence of oligomerization^57^. In contrast, TMCs are predicted to have multiple transmembrane domains and to adopt the topology of other membrane proteins including ion channels and scramblases^58–62^. In addition, TMC1 satisfies the criteria necessary to be the pore-forming subunit of the inner-ear hair-cell mechanotransduction complex. TMC1 is the protein product of a deafness-causing gene in mice and humans^63–65^ and is essential for mouse inner-ear mechanotransduction along with TMC2, a closely related homolog^44,45,66^. Hair cells from *Tmc1*/*Tmc2* double knockout mice lack mechanotransduction currents, and transfection of *Tmc1* or *Tmc2* genes rescues hair-cell mechanotransduction^44^. Both TMC1 and TMC2 proteins are localized to the site of transduction at the base of the tip link and are expressed at the onset of mechanotransduction with TMC1 expression dominating throughout adulthood^44–46,67^. Importantly, several deafness-causing mutations in TMC1, including the “Beethoven” M412K mutation, alter the electrophysiological properties of the hair-cell mechanotransduction channel^45,60,63,68^. In addition, the *Danio rerio tmc1* and *tmc2a*/*tmc2b* orthologs are essential for fish hair-cell mechanotransduction^69^ and electrophysiological experiments using the green sea turtle TMC1 and the budgerigar TMC2 proteins reconstituted in proteoliposomes demonstrated that TMCs can form ion channels gated by membrane tension without any accessory proteins^70^. Together, these experimental results strongly suggest that TMC1 and TMC2 are the pore-forming subunits of the mechanotransduction complex in vertebrate hair cells. Yet the mechanisms of ion permeation and activation still remain poorly defined, in part because of a lack of experimentally derived structural models for TMC proteins and of the complexes they may form with PCDH15, TMHS, and TMIE.

Bioinformatics analyses of TMC protein sequences show that TMCs are evolutionarily related to the TMEM16 family of lipid scramblases and ion channels (also known as anoctamins)^58,61,62^. Structural studies show that TMEM16 proteins form homodimers with each monomer having ten transmembrane domains labeled α1 to α10 ^71–78^. Intriguingly, bioinformatics and structural analyses show that the OSCA/TMEM63 family of mechanosensitive channels have a similar homodimeric architecture with each monomer featuring eleven transmembrane domains, ten of which adopt a TMEM16-like fold^79–82^. Biochemical assays and low-resolution electron microscopy images of purified TMC1 showed that this protein also assembles as a dimer^60^, suggesting that TMC proteins adopt the oligomeric state and folding topology of TMEM16 and OSCA/TMEM63 proteins.

The most striking feature of TMEM16 and OSCA protein structures is their lipid-facing, groove-forming active region in each monomer^71–82^, as opposed to a central cavity in conventional ion channels. Based on sequence and oligomeric state similarities between TMEM16 and TMCs^58,61^, we developed TMEM16-based dimeric comparative models of TMC1 that have a lipid-facing groove forming a putative pore^59,60^. These models were used to successfully explain *ex-vivo* electrophysiological data obtained with hair cells expressing cysteine-substituted TMC1 mutants^60^. In these experiments, hair-cell channel conductivity was altered when cysteine residues predicted to be at the TMC1 pore in our model were modified with blocking methanethiosulfonate (MTS) reagents, thus confirming that these residues were accessible and lined the pore of the hair-cell mechanotransduction complex^60^. In parallel, similar TMEM16-based models were used to identify the putative TMC1 pore and to guide the interpretation of experiments showing TMC1/TMC2-dependent dextran permeation of hair cells^59^. Remarkably, deafness-causing mutations known to alter the permeation properties of the mechanotransduction channel localize to the pore region in the TMEM16-based TMC1 models, further suggesting that TMC proteins are the pore-forming subunits of the mechanotransduction complex and validating our TMC1 models^59,60^.

In previous work, equilibrium molecular dynamics (MD) simulations of our TMEM16-based TMC1 comparative models revealed potassium ions spontaneously exploring the putative pore, even though all models were based on structures of lipid scramblases or anion channels^60^. This solidified the notion that TMC proteins can be cation channels, but left open questions related to whether these models represented open states with the expected conductance for the mechanotransduction channel in the presence of the native hair-cell transmembrane potential. Here we use non-equilibrium MD simulations of human TMC1 models to predict their conductance in the presence of transmembrane potentials. These simulations revealed that some TMEM16-based TMC1 models can conduct cations at rates comparable to those measured for the native mechanotransduction channel and suggest that gating mechanisms might involve both changes in protein conformation and membrane tension.

## RESULTS

### A TMC1 model reveals membrane-exposed and hydrated pores

To run equilibrium and non-equilibrium MD simulations of TMC1 we first created an I-TASSER^83^ comparative model (model 1) based on the structure of the *Nectria haematococca* lipid scramblase TMEM16 (*nh*TMEM16, PDB: 4WIS; Fig. 1a)^71^ and on an alignment between *nh*TMEM16 and human TMC1 sequences following Ballesteros *et al.*^59^. The TMC1 residues probed in the cysteine mutagenesis experiments by Pan *et al.*^60^ that lined the pore of the transduction channel complex in murine hair cells were also found in the pore-forming region of this TMC1 model (Fig. 1b). Then, we built an all-atom molecular simulation system comprising the full-length TMC1 homodimer channel complex with its two pores. This system included N- and C-terminal domains and helices α1 to α10 (Supplementary Alignment) for monomers A and B, embedded in a pure 1-palmitoyl-2-oleoyl-sn-glycero-3-phosphocholine (POPC) bilayer with explicit water at physiological ionic concentration (0.150 M KCl as expected for inner-ear endolymph^84^; Fig. 1a). Although phosphatidylinositol 4,5-bisphosphate (PIP2) and cholesterol are important for hair-cell mechanotransduction^43,85,86^, simulations with a pure POPC membrane are essential to establish a baseline for future studies with heterogeneous lipids. During an initial 100-ns long MD equilibration of the complete all-atom system without transmembrane potential (Sim1a and Sim1b; Table 1) we observed stable water channels at the hydrophilic grooves formed by helices α4 to α7 of each protein monomer (Fig. 1c). Lipid head-groups, initially at the typical upper and lower edges of the bilayer, provided additional stability to these water channels by lining themselves along the groove and forming a hydrophilic wall that separated water from the hydrophobic bilayer core formed by lipid tails. The concentrations of water molecules in the pore of each monomer were comparable, in some places, to bulk water concentrations, suggesting that each monomer may have a solvent-accessible pore able to conduct ions upon application of a transmembrane potential (Fig. 1d).

**Table 1.**
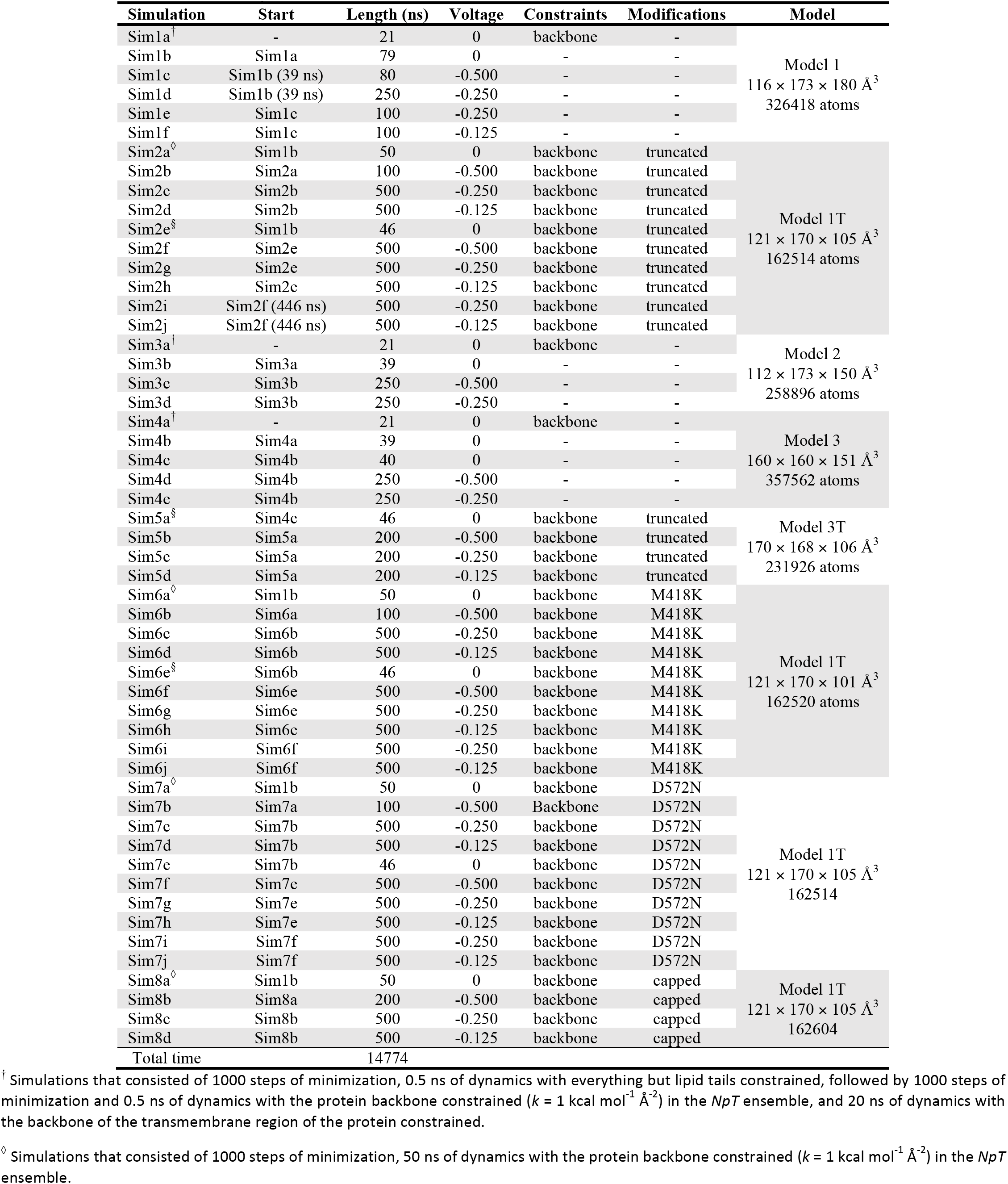
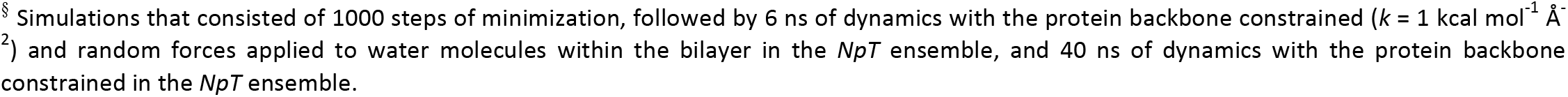
Summary of simulations.

| Simulation | Start | Length (ns) | Voltage | Constraints | Modifications | Model |
| --- | --- | --- | --- | --- | --- | --- |
| Sim1a<sup>†</sup> | - | 21 | 0 | backbone | - | Model 1<br>116 × 173 × 180 Å<sup>3</sup><br>326418 atoms |
| Sim1b | Sim1a | 79 | 0 | - | - |  |
| Sim1c | Sim1b (39 ns) | 80 | -0.500 | - | - |  |
| Sim1d | Sim1b (39 ns) | 250 | -0.250 | - | - |  |
| Sim1e | Sim1c | 100 | -0.250 | - | - |  |
| Sim1f | Sim1c | 100 | -0.125 | - | - |  |
| Sim2a<sup>◇</sup> | Sim1b | 50 | 0 | backbone | truncated | Model 1T<br>121 × 170 × 105 Å<sup>3</sup><br>162514 atoms |
| Sim2b | Sim2a | 100 | -0.500 | backbone | truncated |  |
| Sim2c | Sim2b | 500 | -0.250 | backbone | truncated |  |
| Sim2d | Sim2b | 500 | -0.125 | backbone | truncated |  |
| Sim2e<sup>§</sup> | Sim1b | 46 | 0 | backbone | truncated |  |
| Sim2f | Sim2e | 500 | -0.500 | backbone | truncated |  |
| Sim2g | Sim2e | 500 | -0.250 | backbone | truncated |  |
| Sim2h | Sim2e | 500 | -0.125 | backbone | truncated |  |
| Sim2i | Sim2f (446 ns) | 500 | -0.250 | backbone | truncated |  |
| Sim2j | Sim2f (446 ns) | 500 | -0.125 | backbone | truncated |  |
| Sim3a<sup>†</sup> | - | 21 | 0 | backbone | - | Model 2<br>112 × 173 × 150 Å<sup>3</sup><br>258896 atoms |
| Sim3b | Sim3a | 39 | 0 | - | - |  |
| Sim3c | Sim3b | 250 | -0.500 | - | - |  |
| Sim3d | Sim3b | 250 | -0.250 | - | - |  |
| Sim4a<sup>†</sup> | - | 21 | 0 | backbone | - | Model 3<br>160 × 160 × 151 Å<sup>3</sup><br>357562 atoms |
| Sim4b | Sim4a | 39 | 0 | - | - |  |
| Sim4c | Sim4b | 40 | 0 | - | - |  |
| Sim4d | Sim4b | 250 | -0.500 | - | - |  |
| Sim4e | Sim4b | 250 | -0.250 | - | - |  |
| Sim5a<sup>§</sup> | Sim4c | 46 | 0 | backbone | truncated | Model 3T<br>170 × 168 × 106 Å<sup>3</sup><br>231926 atoms |
| Sim5b | Sim5a | 200 | -0.500 | backbone | truncated |  |
| Sim5c | Sim5a | 200 | -0.250 | backbone | truncated |  |
| Sim5d | Sim5a | 200 | -0.125 | backbone | truncated |  |
| Sim6a<sup>◇</sup> | Sim1b | 50 | 0 | backbone | M418K | Model 1T<br>121 × 170 × 101 Å<sup>3</sup><br>162520 atoms |
| Sim6b | Sim6a | 100 | -0.500 | backbone | M418K |  |
| Sim6c | Sim6b | 500 | -0.250 | backbone | M418K |  |
| Sim6d | Sim6b | 500 | -0.125 | backbone | M418K |  |
| Sim6e<sup>§</sup> | Sim6b | 46 | 0 | backbone | M418K |  |
| Sim6f | Sim6e | 500 | -0.500 | backbone | M418K |  |
| Sim6g | Sim6e | 500 | -0.250 | backbone | M418K |  |
| Sim6h | Sim6e | 500 | -0.125 | backbone | M418K |  |
| Sim6i | Sim6f | 500 | -0.250 | backbone | M418K |  |
| Sim6j | Sim6f | 500 | -0.125 | backbone | M418K |  |
| Sim7a<sup>◇</sup> | Sim1b | 50 | 0 | backbone | D572N | Model 1T<br>121 × 170 × 105 Å<sup>3</sup><br>162514 |
| Sim7b | Sim7a | 100 | -0.500 | Backbone | D572N |  |
| Sim7c | Sim7b | 500 | -0.250 | backbone | D572N |  |
| Sim7d | Sim7b | 500 | -0.125 | backbone | D572N |  |
| Sim7e | Sim7b | 46 | 0 | backbone | D572N |  |
| Sim7f | Sim7e | 500 | -0.500 | backbone | D572N |  |
| Sim7g | Sim7e | 500 | -0.250 | backbone | D572N |  |
| Sim7h | Sim7e | 500 | -0.125 | backbone | D572N |  |
| Sim7i | Sim7f | 500 | -0.250 | backbone | D572N |  |
| Sim7j | Sim7f | 500 | -0.125 | backbone | D572N |  |
| Sim8a<sup>◇</sup> | Sim1b | 50 | 0 | backbone | capped | Model 1T<br>121 × 170 × 105 Å<sup>3</sup><br>162604 |
| Sim8b | Sim8a | 200 | -0.500 | backbone | capped |  |
| Sim8c | Sim8b | 500 | -0.250 | backbone | capped |  |
| Sim8d | Sim8b | 500 | -0.125 | backbone | capped |  |
| Total time |  | 14774 |  |  |  |  |

<sup>†</sup> Simulations that consisted of 1000 steps of minimization, 0.5 ns of dynamics with everything but lipid tails constrained, followed by 1000 steps of minimization and 0.5 ns of dynamics with the protein backbone constrained ( $k = 1 \text{ kcal mol}^{-1} \text{ \AA}^{-2}$ ) in the $NpT$ ensemble, and 20 ns of dynamics with the backbone of the transmembrane region of the protein constrained.
<sup>◇</sup> Simulations that consisted of 1000 steps of minimization, 50 ns of dynamics with the protein backbone constrained ( $k = 1 \text{ kcal mol}^{-1} \text{ \AA}^{-2}$ ) in the $NpT$ ensemble.
§ Simulations that consisted of 1000 steps of minimization, followed by 6 ns of dynamics with the protein backbone constrained ( $k = 1 \text{ kcal mol}^{-1} \text{ \AA}^{-2}$ ) and random forces applied to water molecules within the bilayer in the $NpT$ ensemble, and 40 ns of dynamics with the protein backbone constrained in the $NpT$ ensemble.

**Fig. 1.**
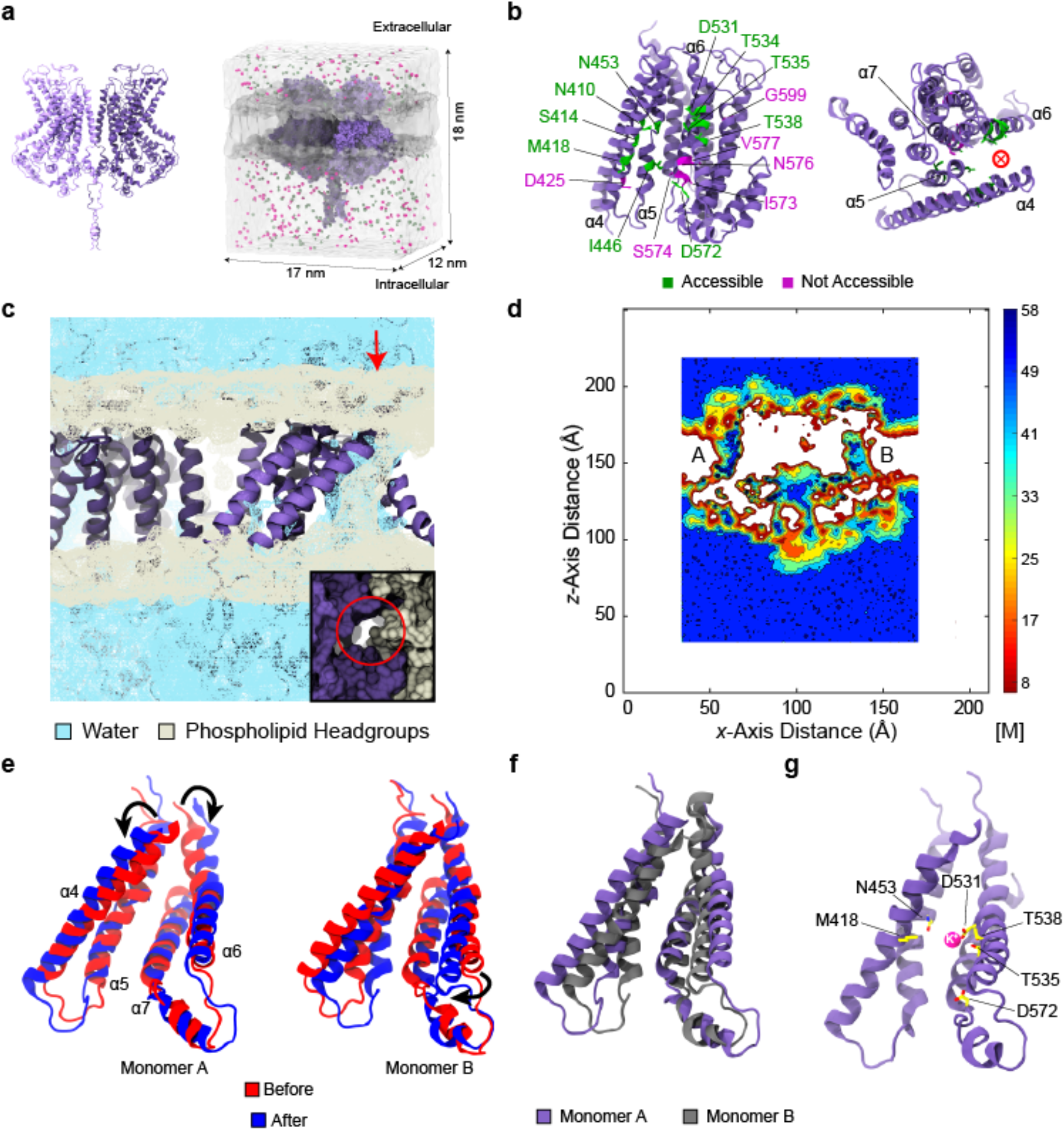
Lipid-exposed pore in a TMEM16-based TMC1 model. **a** A TMC1 dimer built using the *nh*TMEM16 structure as a template (model 1). Simulation system is shown on the right with protein in bright and dark purple colors. Water is in transparent white, lipid membrane in transparent gray, potassium and chloride ions are pink and green, respectively. **b** Side and top views of TMC1 transmembrane helices from model 1 shown as ribbons (monomer). Helices α4 to α7 form the conductive groove facing lipids. Residues probed by Pan *et al.*^60^ are shown as sticks. Solvent exposed residues are shown in green, others are in magenta. Left - front view with pore facing viewer. Right - top view looking down from the extracellular side. **c** Density of phospholipid head groups (light yellow) and water molecules (sky blue) averaged over an equilibrium simulation (Sim1b). Inset shows top view of the pore (red circle). **d** Average water concentration on a plane passing through the protein (60 ns; Sim1b). **e** Relative positions of pore-forming helices before (red) and after (blue) equilibration for monomer A (left) and monomer B (right). **f** TMC1 monomers A (purple) and B (gray) superposed after 60 ns of equilibration (Sim1b). **g** Potassium ion (pink) exploring the pore of monomer B during an equilibrium simulation (Sim1a and Sim1b). Key residues that line the pore are shown.

Further analysis of the equilibrium trajectory without a transmembrane potential (first 60 ns of Sim1a and Sim1b) revealed that the average radius over time (Fig. S1a), the minimum pore radius as a function of time (Fig. S1b), and the time-average of pore radii at different positions across the membrane (Fig. S1c) were similar for monomers A and B. However, two of the pore-forming helices, α4 and α6, had distinct dynamics and adopted different conformations in each TMC1 monomer. In monomer A, both α4 and α6 helices moved away from each other to widen the extracellular opening of the putative pore, whereas the same helices in monomer B moved to occlude the extracellular (α4) and intracellular (α6) sides of the pore (Fig. 1e,f). Consistently, the distance between residues Met^418^ in α4 and Thr^535^ in α6 increased for monomer A and decreased for monomer B during the equilibrium simulation (Fig. S1d). These data suggest that relative movement of helices α4 and α6 control pore aperture and the open-close equilibrium as observed for TMEM16 lipid scramblases^75^. Additionally, the distinct conformational changes observed in each monomer suggest that each pore in the channel complex can adopt different functional states independently.

Interestingly, during equilibration (Sim1a and Sim1b) a potassium ion explored the pore groove of monomer B without fully crossing from one membrane side to the other (Fig. 1g; Movie 1). This occurred spontaneously in the absence of a transmembrane potential, suggesting that the pore in monomer B, and likely the wider and hydrated pore in monomer A, are both in potentially conductive states.

### Large cationic conductance in a full-length TMC1 model

To investigate TMC1 conduction properties we performed simulations in which transmembrane potentials of −0.500 V (Sim1c) and −0.250 V (Sim1d) were applied using an external electric field^87,88^ (Tables 1 and 2). These transmembrane potentials are large compared to physiological values (up to - 0.125 V)^89^, but allowed us to explore conduction events on short simulation timescales, as routinely done in MD simulations of ion channel systems^87,88,90–94^. To determine whether helices α4 and α6 would influence conductance, we chose a TMC1 conformation in which the Met^418^ - Thr^535^ distance was large for monomer A and small for monomer B, obtained after 60 ns of equilibration for the full-length model 1 (Fig. S1d). Thus, simulations at −0.500 V and −0.250 V started from this state. Average electrostatic potential maps confirmed the presence of the transmembrane potential and showed a large potential barrier due to the lipid membrane along with narrow paths with steep potential gradients through the pores (Fig. S2a,b). During both simulations, potassium ions travelled from the extracellular to the intracellular side while a few chloride ions occasionally flowed in the opposite direction (Fig. S3a,b; Movie 2). The potassium ions quickly traveled to the center of the pore formed by α4, α5, α6, and α7 as they moved towards its intracellular side. However, in rare instances, potassium ions entered through the side of α4 facing the lipids, rather than through the primary permeation pathway. Conductance of the TMC1 channel complex during the simulation at −0.500 V was estimated to be between 66 pS and 104 pS and was mainly driven by potassium ions flowing through monomer A (80 ns; 26 permeation events total; lower estimate obtained from scaling as described in Table 2 and Methods; Fig. S3a). These data are consistent with a trend observed for average pore radii as a function of time, which were often larger for the pore in monomer A (Fig. S3d).

**Table 2.** Ion conduction^†^.

| Label | Length (ns) | Voltage | K<sup>+</sup><sub>+z</sub> | K<sup>+</sup><sub>-z</sub> | Cl<sup>-</sup><sub>+z</sub> | Cl<sup>-</sup><sub>-z</sub> | I pA | C (pS) |
| --- | --- | --- | --- | --- | --- | --- | --- | --- |
| Sim1a | 21 | 0 | 0 | 0 | 0 | 0 | 0 | 0 |
| Sim1b | 79 | 0 | 0 | 0 | 0 | 0 | 0 | 0 |
| Sim1c | 80 | -0.500 | 0 | 23 | 3 | 0 | 52.07 | 104.13 |
| Sim1d | 250 | -0.250 | 0 | 6 | 1 | 0 | 4.49 | 17.94 |
| Sim1e | 100 | -0.250 | 0 | 11 | 2 | 0 | 20.83 | 83.30 |
| Sim1f | 100 | -0.125 | 0 | 2 | 1 | 0 | 4.81 | 38.45 |
| Sim2a | 50 | 0 | 0 | 0 | 0 | 0 | 0 | 0 |
| Sim2b | 100 | -0.500 | 0 | 39 | 8 | 0 | 75.29 | 150.59 |
| Sim2c | 500 | -0.250 | 0 | 29 | 3 | 0 | 10.25 | 41.01 |
| Sim2d | 500 | -0.125 | 0 | 8 | 0 | 0 | 2.56 | 25.51 |
| Sim2e | 46 | 0 | 0 | 0 | 0 | 0 | 0 | 0 |
| Sim2f | 500 | -0.500 | 0 | 216 | 83 | 0 | 95.80 | 191.60 |
| Sim2g | 500 | -0.250 | 0 | 28 | 4 | 0 | 10.25 | 41.01 |
| Sim2h | 500 | -0.125 | 0 | 4 | 1 | 0 | 1.60 | 12.82 |
| Sim2i | 500 | -0.250 | 0 | 39 | 3 | 0 | 13.46 | 53.83 |
| Sim2j | 500 | -0.125 | 0 | 17 | 2 | 0 | 6.09 | 48.70 |
| Sim3a | 21 | 0 | 0 | 0 | 0 | 0 | 0 | 0 |
| Sim3b | 39 | 0 | 0 | 0 | 0 | 0 | 0 | 0 |
| Sim3c | 250 | -0.500 | 0 | 106 | 184 | 0 | 185.83 | 371.66 |
| Sim3d | 250 | -0.250 | 0 | 2 | 0 | 0 | 1.28 | 5.13 |
| Sim4a | 21 | 0 | 0 | 0 | 0 | 0 | 0 | 0 |
| Sim4b | 39 | 0 | 0 | 0 | 0 | 0 | 0 | 0 |
| Sim4c | 40 | 0 | 0 | 0 | 0 | 0 | 0 | 0 |
| Sim4d | 250 | -0.500 | 0 | 2 | 1 | 0 | 1.92 | 3.84 |
| Sim4e | 250 | -0.250 | 0 | 1 | 2 | 0 | 1.92 | 7.69 |
| Sim5a | 46 | 0 | 0 | 0 | 0 | 0 | 0 | 0 |
| Sim5b | 200 | -0.500 | 0 | 0 | 0 | 0 | 0 | 0 |
| Sim5c | 200 | -0.250 | 0 | 0 | 0 | 0 | 0 | 0 |
| Sim5d | 200 | -0.125 | 0 | 0 | 0 | 0 | 0 | 0 |
| Sim6a | 50 | 0 | 0 | 0 | 0 | 0 | 0 | 0 |
| Sim6b | 100 | -0.500 | 0 | 53 | 23 | 0 | 121.75 | 243.50 |
| Sim6c | 500 | -0.250 | 0 | 32 | 13 | 0 | 14.42 | 57.67 |
| Sim6d | 500 | -0.125 | 0 | 9 | 6 | 0 | 4.81 | 38.45 |
| Sim6e | 46 | 0 | 0 | 0 | 0 | 0 | 0 | 0 |
| Sim6f | 500 | -0.500 | 0 | 258 | 310 | 0 | 181.99 | 363.97 |
| Sim6g | 500 | -0.250 | 0 | 34 | 45 | 0 | 25.31 | 101.25 |
| Sim6h | 500 | -0.125 | 0 | 8 | 5 | 0 | 4.17 | 33.28 |
| Sim6i | 500 | -0.250 | 0 | 38 | 30 | 0 | 21.79 | 87.15 |
| Sim6j | 500 | -0.125 | 0 | 4 | 4 | 0 | 2.56 | 20.51 |
| Sim7a | 50 | 0 | 0 | 0 | 0 | 0 | 0 | 0 |
| Sim7b | 100 | -0.500 | 0 | 23 | 15 | 0 | 60.88 | 121.75 |
| Sim7c | 500 | -0.250 | 0 | 21 | 7 | 0 | 8.97 | 35.88 |
| Sim7d | 500 | -0.125 | 0 | 8 | 9 | 0 | 5.45 | 43.57 |
| Sim7e | 46 | 0 | 0 | 0 | 0 | 0 | 0 | 0 |
| Sim7f | 500 | -0.500 | 0 | 156 | 81 | 0 | 75.93 | 151.87 |
| Sim7g | 500 | -0.250 | 0 | 5 | 1 | 0 | 1.92 | 7.69 |
| Sim7h | 500 | -0.125 | 0 | 6 | 1 | 0 | 2.24 | 17.94 |
| Sim7i | 500 | -0.250 | 0 | 40 | 25 | 0 | 20.83 | 83.30 |
| Sim7j | 500 | -0.125 | 0 | 2 | 1 | 0 | 0.96 | 7.69 |
| Sim8a | 50 | 0 | 0 | 0 | 0 | 0 | 0 | 0 |
| Sim8b | 200 | -0.500 | 0 | 77 | 5 | 0 | 65.68 | 131.36 |
| Sim8c | 500 | -0.250 | 0 | 38 | 2 | 0 | 12.82 | 51.26 |
| Sim8d | 500 | -0.125 | 0 | 8 | 2 | 0 | 3.20 | 25.63 |

<sup>†</sup> Events, currents and conductance values are reported for combined monomers A and B.

Similarly, time-averaged radii at the intracellular entrance of the pore and the Met^418^ - Thr^535^ distance as a function of time were larger for monomer A (Fig. S3g,j). In contrast, conductance of the TMC1 channel complex during the simulation at - 0.250 V was between 11 pS and 18 pS (250 ns; 7 permeation events; Fig. S3b), indicating a lower albeit not negligible current driven by potassium ions flowing through pores of both monomers. Consistently, the average pore radii for monomer A at −0.250 V decreased while the average pore radii for monomer B remained at values similar to those observed in the simulation carried out at −0.500 V (cf. Fig. S3e and Fig. S3d). Similarly, the Met^418^ - Thr^535^ distance as a function of time decreased and the time-averaged radii at the intracellular entrance of the pore was similar for both monomers (Fig. S3h,k). These results indicate that the pore of monomer A in our model fluctuates between two conformational states, one with high conductance and one with lower conductance, while the pore of monomer B stayed in a conformation with low conductance at the transmembrane potentials tested.

Next, we asked whether the high-conductance state observed for our TMC1 model simulated at −0.500 V would remain conductive at lower transmembrane potentials, namely −0.250 V (Sim1e) and −0.125 V (Sim1f). At the end of the simulation done at −0.500 V (Sim1c), we instantly decreased the voltage *in silico* from −0.500 V to −0.250 V and to −0.125 V to continue two separate simulations in parallel for 100 ns each. Average electrostatic potential maps showed narrow paths with steep potential gradients through the pores (Fig. 2a and S2c). For the simulation done with a ramped-down transmembrane potential of −0.250 V (Sim1e) we observed that both pores conducted several potassium ions and recorded a conductance for the TMC1 complex between 53 pS and 83 pS (100 ns; 13 permeation events; Fig. 2b), with monomer A carrying most of the ionic current as observed at −0.500 V. Interestingly, two chloride ions (one per pore) permeated to contribute to this conductance. Trends for average radii over time (Fig. 2c) and for the time-averaged radii at the intracellular entrance (Fig. 2d) correlated with the conduction state of each monomer, as did the Met^418^ - Thr^535^ distance (Fig. 2e). During the simulation done with a ramped-down transmembrane potential of −0.125 V (Sim1f) we observed only two permeation events of potassium and one of chloride ions, with an estimated conductance for the TMC1 complex between 25 pS and 39 pS (100 ns; 3 permeation events; Fig. S2c and S3c). The average radii over time and the time-averaged radius across the membrane were similar among the pores in both monomers (Fig. S3f,i), but in this case the Met^418^ - Thr^535^ distance remained larger for pore A than for pore B (Fig. S3l), suggesting that conformational dynamics of lipids also contribute to determine the conductive state of each pore.

**Fig. 2.**
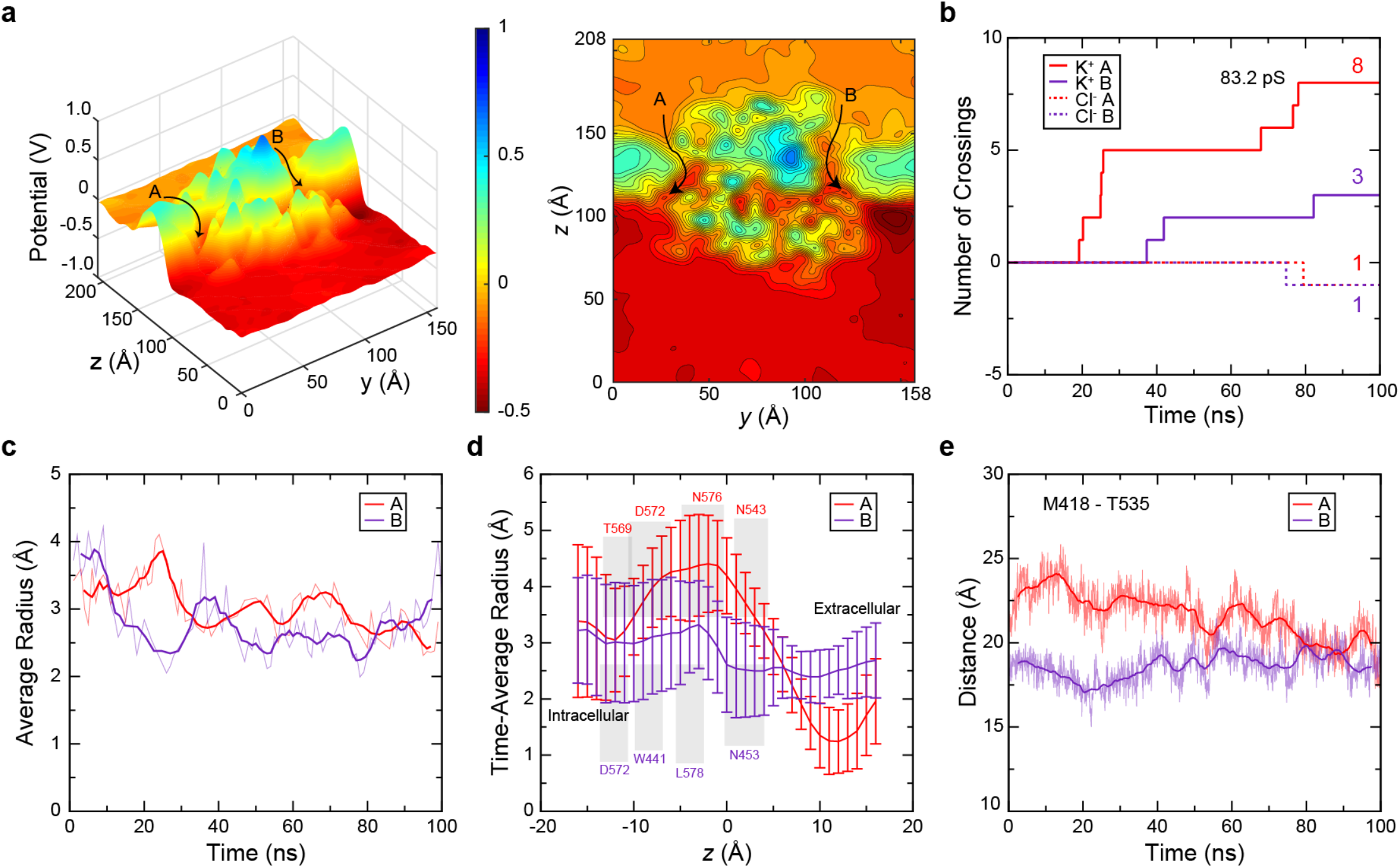
Ion conduction and pore properties of TMC1. **a** Left – Cross-section slice through the transmembrane pores showing the electrostatic potential surface averaged over 100 ns of simulation of the TMC1 model at −0.250 V (Sim1e). Probable ion permeation pathways shown by arrows. Right – A two-dimensional contour plot of the same potential surface. **b** Number of ion crossings as a function of time (Sim1e). Solid lines are for potassium ions going through monomers A (red) and B (indigo). Dashed lines are for chloride ions going through monomers A (red) and B (indigo). **c** Average radius across the pore axis (*z*) for each monomer as a function of time (Sim1e). Transparent lines show raw data after curation (see Methods) while solid lines indicate their 5-ns running averages. **d** Time-averaged radii as a function of position (*z*) along the pore axis. Value of *z* increases from intra- to extra-cellular sides. Error bars are standard deviation. Key residues identified in the HOLE output are shown. **e** Separation between C_α_ atoms of Met^418^ in α4 and Thr^535^ in α6. Transparent lines indicate raw data recorded every 50 ps, solid lines are 1 ns running averages.

A detailed analysis of the trajectory for the simulations of full-length TMC1 (model 1) in which a transmembrane potential of −0.500 V was applied (Sim1c) revealed a patch of hydrophobic residues that often interacted with lipids, including Phe^363^, Phe^366^, Leu^367^, Met^411^, Val^412^, Met^418^, Phe^419^, Leu^533^, Ile^541^, and Phe^544^ (Fig. S4a,b). These residues might be essential for TMC1 embedding in a bilayer and may sense changes in membrane properties and tension regulating the opening and closure of the pore when the membrane is stretched. An analysis of monomer A and its pore during trajectories for simulations with transmembrane potential (Sim1c through Sim1f) revealed pore-lining residues (Fig. S4c), including several that interacted with potassium ions at the extracellular entrance (Glu^405^, Glu^464^, Glu^523^), middle (Asn^453^, Asp^531^), and at the intracellular exit of the channel (Asp^572^). Lipids lined most of the intracellular side of the transmembrane pore (Fig. S4c), thereby supporting the dual protein-lipid nature of the TMC1 channel.

### Large cationic conductance in a minimalistic TMC1 model

To predict ion conduction properties of TMC1 at longer simulation timescales, we constructed a minimal molecular simulation system with a truncated version of the TMC1 dimer consisting of the transmembrane helices and some loops of the protein (model 1T), all with their backbones constrained (Fig. 3a). Given that sequence similarity between TMC1 and *nh*TMEM16 is highest for transmembrane domains, this model had the most reliable structural predictions at the cost of missing backbone dynamics and some intra- and extra-cellular loops. To capture the high-conductance state described above, we chose a protein conformation in which the Met^418^ - Thr^535^ distance was large for both monomers A and B, obtained after a 100 ns equilibration of the full-length TMC1 model 1 (Fig. S1d). Equilibration of the truncated model (Sim2a) was followed by a simulation with a transmembrane potential of −0.500 V (100 ns; Sim2b), which when completed was used as a starting point for two independent simulations in which the voltage was instantly decreased to −0.250 V (500 ns; Sim2c; Fig. 3b-d, S6a, and S7b) or to −0.125 V (500 ns; Sim2d). Conductance of the truncated TMC1 channel complex during the simulation at - 0.500 V was estimated to be between 95 pS and 151 pS (100 ns; 47 permeation events) and was mainly driven by potassium ions flowing through monomer A (Fig. S5a), although there were 8 chloride ions contributing to the ionic current. Conductance of the TMC1 channel complex during the simulation at −0.250 V was estimated to be between 26 pS and 41 pS (500 ns; 32 permeation events; three chloride ions; Figs. 3d, S6a and S7b), and for the simulation at −0.125 V was between 16 pS and 26 pS (500 ns; 8 permeation events; cero chloride ions; Fig. S5d). These results indicate that eliminating intra- and extra-cellular loops increases TMC1 conductance and decreases selectivity for potassium versus chloride at a high transmembrane potential, although conduction of chloride ions remains small. The voltage-dependence of conductance observed in our backbone-constrained minimalistic TMC1 model suggests that residue side chains and lipids can accommodate states of variable conductance with wider pores (Fig. S7), likely because forced passage of ions with their hydration shells driven by steep electrostatic potential drops push away the malleable lipid wall present in our TMC1 model.

**Fig. 3.**
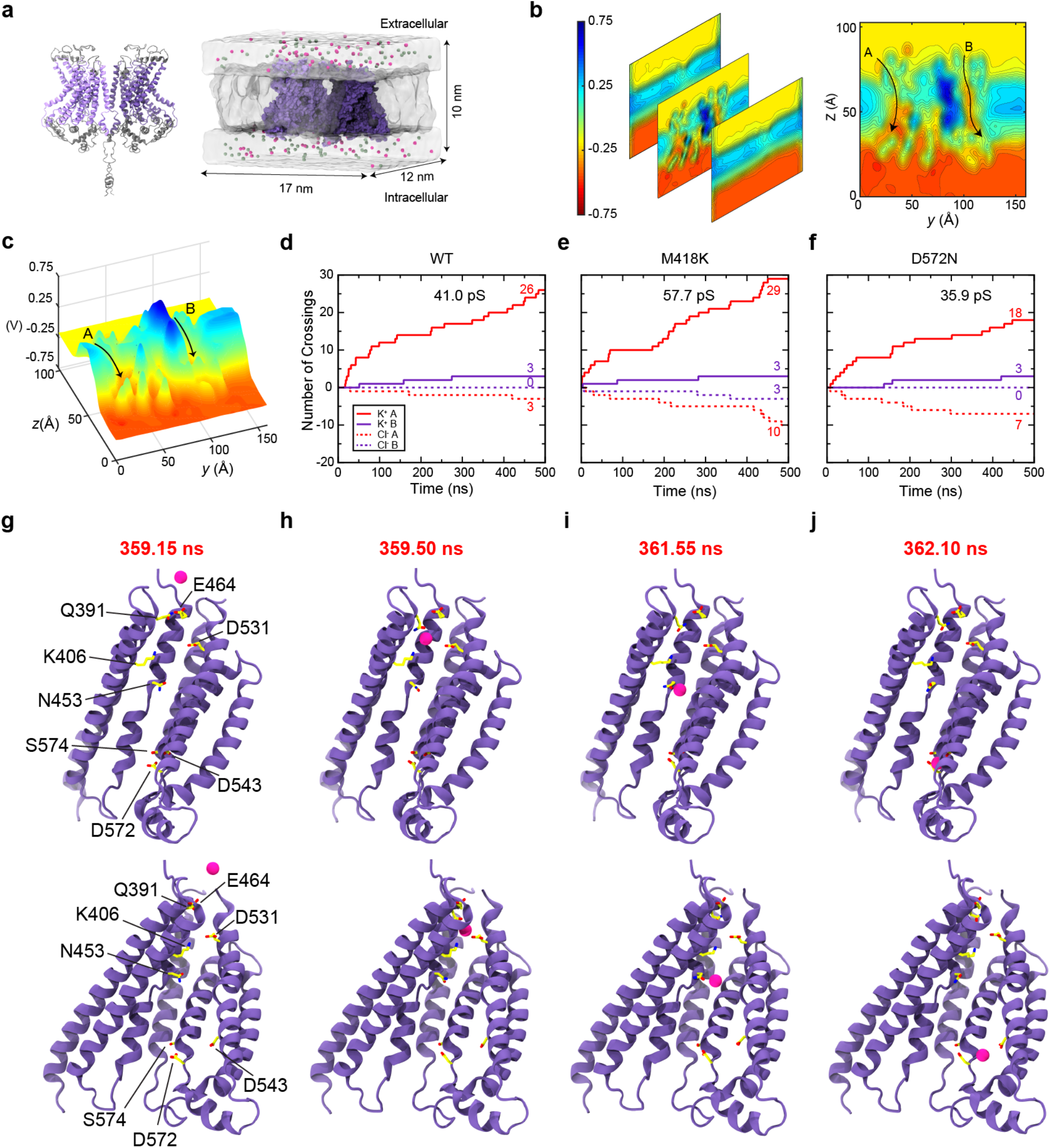
Ion conduction through a minimalistic truncated model of the TMC1 dimer. **a** Left – truncation scheme used to create the minimalistic all atom simulation system. Excised intra- and extra-cellular regions are depicted in gray. On the right – the complete simulation system of the minimalistic model shown in surface representation. Protein is shown in bright and dark purple colors, water is shown in transparent white, lipid membrane is shown in transparent gray, potassium and chloride ions are shown as pink and green spheres, respectively. **b** Two-dimensional contour plots of the electrostatic potential surface averaged over 500 ns for the minimalistic TMC1 model at −0.250 V (Sim2c). Left – Each cross section corresponds to a *y-z* slice perpendicular to the membrane plane at three different points along the *x* axis from back to front: a point behind the protein dimer, a point cutting through the TMC1, and a point in front of the dimer. Right – Middle cross section shown as a flat surface. Black arrows show pore regions. **c** A three-dimensional representation of the two-dimensional plot in **b**. **d-f** Ion crossing events through the pore of the wild type, M418K, and D572N truncated TMC1 systems at −0.250 V (Sim2c, Sim6c, and Sim7c). Solid and dashed lines denote potassium and chloride permeation events, respectively. Also shown in Fig. S6a-c. **g-j** Snapshots showing a potassium permeation event during a simulation of the truncated wild-type TMC1 model at −0.250 V (Sim2c). Residues that were within 3 Å of the ion are shown in sticks and labeled. Top and bottom rows show two different views at identical times.

Analysis of ion permeation during simulations of the truncated TMC1 complex at −0.250 V (500 ns; 32 permeation events; three chloride ions; Figs. 3d and S6a) revealed rapid passage of potassium ions, which interacted with several residues that lined the pore. An example of a permeation event (Fig. 3g-j) revealed that a potassium ion was within 3 Å of residues that also lined the pore of the full-length TMC1 channel complex (model 1; Fig. S4c), including Q391, K406, N453, E464, D531, and D572. Interactions with residues D543 and S574 were also observed. The entire permeation event lasted less than 3 ns (Fig. 4g-j).

**Fig. 4.**
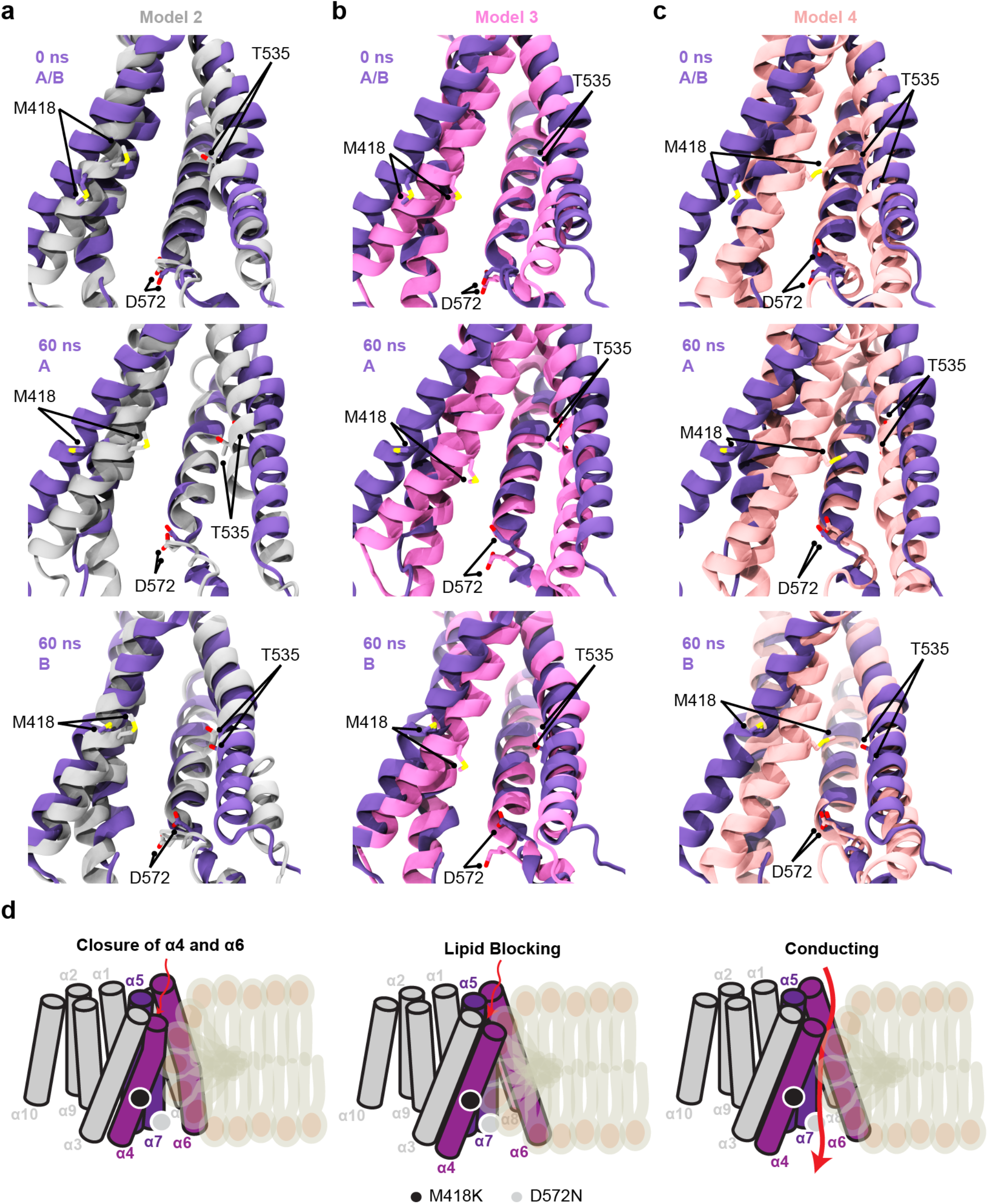
Comparison of TMC1 models. **a** A comparative model of TMC1 based on *nh*TMEM16 (model 1) in purple before (top) and after 60 ns of equilibration (monomers A and B in the middle and bottom panels) overlaid with model 2 in gray. Met^418^, Thr^535^, and Asp^572^ are shown in licorice. **b** Model 1 shown as in **a** overlaid with a comparative model based on *mm*TMEM16A (model 3) in pink. **c** Model 1 shown as in **a** overlaid with an AlphaFold TMC1 model. **d** Schematics of three different states for TMC1. Left – Pore in a closed state due to movement of α4 and α6 to occlude the pore. Middle – Pore widens as helices α4 and α6 separate from each other and become straighter, but lipid head groups block the pore. Right – Conducting state where α4 and α6 are shifted to open the pore and lipid head groups line the pore without blocking it.

To determine whether our results with the TMC1 truncated model (model 1T) were robust or dependent on the state of the lipid bilayer, we did replicate simulations with a system that included the same protein model surrounded by POPC lipids in a different conformation, as these were artificially moved during the first six nanoseconds of a 46-ns long equilibration (Sim2e). The resulting equilibrated system was used as a starting point for 500-ns-long simulations with transmembrane potentials set to −0.500 V (Sim2f), −0.250 V (Sim2g), and −0.125 V (Sim2h). The conductance at a high transmembrane potential was large, between 121 pS and 192 pS, and mainly driven by potassium ions crossing through monomer A, but with a significant number of chloride permeation events as well (299 permeation events; 83 chloride ions). Simulations at reduced transmembrane potentials showed lower conductance values (26 pS to 42 pS at −0.250 V and 8 pS to 13 pS at - 0.125V), consistent with our previous results. We also did simulations that started from a conformation obtained after 446 ns of dynamics of the truncated TMC1 model at −0.500 V (Sim2f) and with voltage instantly decreased to −0.250 V (500 ns; Sim2i) and to −0.125 V (500 ns; Sim2j). Conductance was similar for both ramped-down simulations with estimates between 34 pS and 54 pS at −0.250 V and between 31 pS and 49 pS at −0.125 V. These data suggest that cation conductance is robust with little dependence on the initial lipid configuration around the protein.

Given that truncated models had charged N- and C-termini for each helix, we also created a truncated model of the TMC1 protein where N- and C-termini were neutralized by patching them with acetyl and N-methyl groups, respectively. After a 50-ns long equilibration (Sim8a), we set the transmembrane potential to −0.500 V for 200 ns (Sim8b) and observed permeation of potassium ions with a minor contribution from chloride resulting in an estimated conductance between 83 pS and 131 pS. As the simulation progressed with decreased transmembrane potentials at −0.250 mV (500 ns; Sim8c) or - 0.125 mV (500 ns; Sim8d), the conductance decreased to values between 33 pS and 51 pS, and between 16 pS and 26 pS, respectively. The conductance values estimated for the truncated TMC1 channel complex with and without neutralization of the charged terminal residues are comparable, suggesting that the presence or absence of neutral patched termini did not interfere with ion conduction.

### Helices α4 and α6 in open and closed TMC1 models

Our simulations of full-length and truncated TMC1 dimers (model 1 and model 1T based on *nh*TMEM16) suggest that TMEM16-based models are capable of cation conduction at the rates expected for the hair-cell transduction channel, even in the absence of other transmembrane proteins that may be part of the complex. However, similar simulations of a full-length TMC1 model based on *nh*TMEM16 but using a different sequence alignment (model 2) did not produce consistent results. Equilibration (60 ns; Sim3a and Sim3b) followed by 250 ns of dynamics with a transmembrane potential of −0.500 V (Sim3c) showed an unstable system with rupture of the lipid wall at the dimeric interface and electroporation of ions resulting in an unrealistic high conductance. Conversely, there were only two permeation events for a simulation at a lower transmembrane potential (−0.250 V; Sim3d). A third model based on the structure of the mouse calcium-activated chloride channel TMEM16A (*mm*TMEM16A; PDB: 5OYB)^77^ appeared to be in a closed conformation, as corroborated by simulations of full-length (model 3) and truncated (model 3T) versions that did not show ion conduction in the presence of transmembrane potentials (Sim4a to Sim4e and Sim5a to Sim5d). The main difference between these models was in the relative position of α4, shifted toward the extracellular side in model 2 (Fig. 4a) and displaced to occlude the pore in models 3 and 3T (Fig. 4b). A similar model obtained using AlphaFold 2 ^95,96^, which was not simulated here, also seems to show a closed state (Fig. 4c). These results are consistent with those from our equilibrium simulations of model 1 in which movement of helices α4 and α6 away from each other led to pores with two different conducting states.

### Deafness-causing mutations alter pore electrostatics

Given the success of our TMC1 simulations in reproducing cationic currents at rates that are comparable to those measured for the hair-cell mechanotransduction channel, we decided to use the truncated version (model 1T) to determine the effect that well-characterized deafness-causing TMC1 missense mutations may have on ionic currents. The TMC1 mutations M418K (mouse M412K)^45,63,68,97^ and D572N (mouse D569N)^64,98–101^ cause deafness in humans and mice, and both have been shown to alter resting open probability and calcium permeability, likely without affecting single channel conductance^45,68,101^. Each mutation was introduced to both monomers of the truncated model to create two mutant systems, model 1T M418K and model 1T D572N. Interestingly, simulations of these systems at high transmembrane potentials (−0.500 mV; Sim6b, Sim6f, Sim7b, and Sim7f; Fig. S5b,c) showed large overall conductance caused by increased flow of chloride ions. Similar trends were observed at −0.250 V (Fig. 3d-f), with an average maximum (not scaled, see Methods) conductance of 82 ± 23 pS for the M418K mutant (Sim6c, Sim6g, Sim6i) and of 42 ± 38 pS for the D572N mutant (Sim7c, Sim7g, Sim7i), compared to 45 ± 7 pS for the wild-type protein (Sim2c, Sim2g, Sim2i). The large standard deviation for the D572N system reflects one simulation in which the TMC1 complex conductance was very small (Sim2g). Despite poorer sampling over the 500-ns timescale used at a more physiological transmembrane potential, our data at −0.125 V (Fig. S5d-f) followed trends similar to those observed at −0.250 V, with an average maximum conductance of 31 ± 9 pS for the M418K mutant (Sim6d, Sim6h, Sim6j) and of 23 ± 19 pS for the D572N mutant (Sim7d, Sim7h, Sim7j), compared to 29 ± 18 pS for the wild-type protein (Sim2d, Sim2h, Sim2j). Notably, permeation events through monomer B remained very low, even though mutations were introduced in both monomers. These results suggest that the deafness-causing mutations M418K and D572N do not dramatically impair ionic currents but do change the electrostatics of the pore, which may affect calcium permeability as expected (Fig. S8). Simulations of unconstrained full-length TMC1 models over longer timescales in heterogeneous membranes containing negatively charged lipids such as PIP2 might be necessary to fully capture the effect of these missense mutations on the dynamics and conductance of the hair-cell mechanotransduction channel.

## DISCUSSION

Overall, our simulations of various wild-type and mutant TMC1 models indicate that transmembrane helices α4, α5, α6, and α7, along with lipids surrounding these helices, form the cation-selective pore of TMC1, consistent with previous data^59,60^. We observed that residues at helices α4, α5, α6, and α7 are water and ion accessible, and that one of our TMC1 models is capable of cation conduction at rates expected for the hair-cell mechanotransduction channel, further supporting the presence of a pore in TMC1 and the function of this protein as the mechanotransduction channel of inner-ear hair cells. In our model where TMC1 is embedded on a membrane, lipid head groups spontaneously lined the hydrophilic regions around the pore-forming helices of TMC1, building up a non-proteinaceous wall. Our simulations of these models with applied transmembrane potentials suggest that each monomer can be in one of three functional states: A closed state where α4 and α6 move closer together on the extracellular side (Fig. 4d, left panel); a lipid-occluding state where the conformation of the protein is favorable for pore formation, but lipid head groups are embedded deep inside the pore thereby blocking the permeation pathway (Fig. 4d, middle panel); and a third conducting state where protein and lipids arrange to form a clear pathway for ion conduction (Fig. 4d – right panel). Cation-driven currents were observed for all TMC1 wild-type systems where a protein-lipid pore was in an open state (model 1 and model 1T), with some minor chloride component that increased in the presence of mutations M418K and D572N that altered the electrostatics of the pore.

Residues M418 and D572 face the pore and locate in the middle of α4 and at the intracellular side of α7 of our TMC1 models. Mutation of these residues causes deafness in humans and mice, and electrophysiological characterization of murine hair cells carrying the equivalent TMC1 M412K and D569N mutations shows that these mutations decrease permeability of the mechanotransduction channel to calcium versus cesium^45,60,68,101^. Our simulation results show altered TMC1 pore electrostatics and are consistent with experimental results. Interestingly, M418 is part of a hydrophobic patch of residues that often interacts with lipids, suggesting that this residue not only contributes to ion permeation but could also stabilize TMC1 in the membrane and sense changes in membrane tension. Recent work has shown that the D572N mutation impairs the interaction between TMC1 and TMHS^102^, suggesting an additional role for residue D572 as well. However, it is difficult envision how residue D572 at the intracellular side of the pore would directly interact with TMHS in our TMEM16-based and AlphaFold 2 models of TMC1. Perhaps the effect of the D572N mutation is indirect and relates to changes in conformation or accessibility of adjacent protein regions. Both mutations, M418K and D572N, need to be studied using full-length unconstrained models of TMC1 to fully understand their effects, but our results showing altered pore electrostatics are already consistent with decreased permeability to calcium.

The lipid bilayer plays an important role in gating of several mechanosensitive channels^103–107^, and it is implicated in modulating hair-cell mechanotransduction and adaptation^86,108–110^. Our models predict a tight coupling between the TMC1 pore and membrane lipids, which include PIP2 and rigidifying cholesterol. The negatively charged head groups of PIP2 might directly regulate the nature of the ionic current through the TMC1 pore by blocking the entrance of chloride ions at the intracellular side and by further favoring cation conduction. Similarly, cholesterol, which reduces hair-cell membrane fluidity and restricts channel opening^111^, may limit the movement of lipids around and away from TMC1 preventing its opening and regulating mechanotransduction currents. A dynamic protein-lipid pore, like the one predicted here for TMC1, might also allow for the passage of large cationic molecules through its pore as lipids move away to accommodate them. This would be consistent with experimental results showing that FM1-43, dextrans, and aminoglycosides can permeate the hair-cell transduction channel^59,112–114^.

Our simulations are limited by approximations in all-atom models and force-fields^115,116^, by simulated timescales, and by the lack of an experimental structure of a TMC protein, alone or in complex with potential protein partners. However, our simulation predictions provide strong support for a model in which TMC1 by itself can form a mechanosensitive ion channel gated by membrane tension, in agreement with data obtained with purified TMC1 and TMC2 reconstituted in liposomes^70^, and as observed for bacterial channels^103^ and the structurally similar OSCA/TMEM163 proteins^80^. It is possible that direct coupling to TMIE and TMHS, as well as to calcium- and integrin-binding (CIB) cytoplasmic protein partners is required to tune hair-cell mechanosensitivity or to have a fully functional hair-cell mechanotransduction apparatus^43,49,117,118^. Perhaps TMIE and TMHS or other yet to be identified component of the mechanotransduction complex replaces the lipid wall observed in our simulations of TMC1 alone. It is also feasible that TMC1 and TMC2 just need to be in the neighborhood of a complex formed by the tip link, TMHS, and TMIE, to feel membrane tension upon stretching of PCDH15 ^119^. Although interactions between TMC1/TMC2 and the cytoplasmic domain of PCDH15^120^ suggest a direct coupling of the TMC proteins with the tip link that transfers mechanical stimuli to the mechanotransduction channels, the estimated number of TMC proteins at the stereocilia tip is larger than the number of tip links^24^. This suggests the presence of additional TMC1 and TMC2 proteins that may be able to feel membrane tension upon stretching of PCDH15. In this model, tonotopic conductance changes would not be limited by the discrete number of channels bound per tip link, but rather could have a continuous change modulated by a variable number of TMC channels around the tip link and by the membrane composition^24,86^. Refinement and tests of these predictions will be essential to distinguish between models in which tip links directly pull on the transduction complex or rather communicate with TMC channels through membrane tension.

## METHODS

### Simulated Systems

Four models of human TMC1 were used for analyses. To construct the first TMC1 comparative model, the alignment of the human TMC1 and the *nh*TMEM16 lipid scramblase sequences, as published in Ballesteros *et al.*^59^, was submitted to I-TASSER (Iterative Threading ASSEmbly Refinement) a protein structure prediction web server^121,122^. The *nh*TMEM16 structure (PDB: 4WIS)^71^ was set as a template, and the resulting monomeric model was similar to I-TASSER models that used default sequence alignments. Using VMD^123^, a dimeric model of TMC1 was constructed by aligning the predicted monomeric structure of TMC1 generated by I-TASSER with the monomers of the *nh*TMEM16 structure (PDB: 4WIS)^71^. Clashes between α10 helices from monomers A and B were resolved in COOT^124^. Monomers were moved apart from each other by 4 Å in VMD to further eliminate remaining clashes. This dimer was then minimized for 1000 steps in vacuum using NAMD^125^ and the CHARMM36 force field^126,127^ before embedding it in a POPC lipid bilayer using VMD and its membrane builder plugin. The protein-lipid system was then solvated using TIP3P explicit water and neutralized with 0.150 M KCl to mimic the salt concentration of the endolymph^84^. The atomic coordinates of the full-length system after Sim1b were used to construct a minimal TMC1 system (model 1T, Fig. 2a) with transmembrane domains including residues Lys^151^ to Tyr^215^ (α1), Tyr^252^ to Gly^305^ (α2), Gln^353^ to Ile^469^ (α3 to α5), Trp^516^ to Pro^659^ (α6 to α9), and Pro^686^ to Leu^730^ (α10). The equilibrated lipids from the full-length model were part of the truncated system, which was then solvated and neutralized with 0.150 M KCl. Harmonic constraints (*k* = 1 kcal mol^−1^ Å^−2^) were applied to the entire protein backbone to hold helices in place. The system was minimized for 2000 steps prior to equilibrium and non-equilibrium production runs. Replicates with scrambled lipids were obtained by applying forces to water molecules within the lipid bilayer during the first six nanoseconds of equilibration (Sim2e). The VMD mutator plugin was utilized on the truncated system to construct the mutant truncated TMC1 D572N and M418K systems. Counter ions were added to neutralize systems after mutation, as required for electrostatic computations using the particle mesh Ewald (PME) algorithm.

A second comparative model of TMC1 (model 2) was created using I-TASSER and an alignment of the human TMC1 and the *nh*TMEM16 lipid scramblase sequences as published in Pan *et al.*^60^, with the *nh*TMEM16 structure set as a template (PDB: 4WIS)^71^. The I-TASSER prediction for the monomer was used to build a dimer as described above. Clashes between α10 helices from monomers A and B were resolved in COOT^124^, but monomers were not further separated from each other as done for model 1. The dimer was minimized in vacuum, embedded in a POPC lipid bilayer, and hydrated as done with model 1.

The third model used for simulations, first reported in Pan *et al.*^60^, was generated by submitting to I-TASSER the human TMC1 sequence along with the structure of mouse TMEM16A (PDB: 5OYB)^77^. A dimer was constructed by aligning the I-TASSER generated monomeric model to each monomer of the TMEM16A dimer. This dimer, which had no clashes, was then embedded in a POPC bilayer and solvated, as done for models 1 and 2. The atomic coordinates of the full-length system after Sim4c were used to construct a minimal TMC1 system (model 3T) with transmembrane domains including residues Glu^171^ to Tyr^221^ (α1), Thr^265^ to Asn^309^ (α2), Ser^337^ to Glu^470^ (α3 to α5), Pro^510^ to Ser^661^ (α6 to α9), and Leu^701^ to Asn^745^ (α10). The equilibrated lipids from the full-length model were also used to construct the system, which was then solvated and neutralized (0.150 M KCl). Harmonic constraints (*k* = 1 kcal mol^−1^ Å^−2^) were applied to the entire protein backbone to hold helices in place. The system was minimized for 2000 steps prior to equilibrium and non-equilibrium production runs. The fourth model of human TMC1 was obtained from the AlphaFold Protein Structure Database (Q8TDI8)^95,96^. Size of simulation box for systems including models 1, 1T, 2, 3, and 3T are in Table 1.

### Simulations

All our all-atom MD simulations^90,92,93,115,128–134^ were performed using NAMD 2.12 or 2.13 and the CHARMM36 force field with CMAP corrections^126,127,135^. A 12-Å cutoff distance was employed with a force-based switching function starting at 10 Å. Periodic boundary conditions and the PME method were used to calculate long-range electrostatic interactions with a grid density greater than 1 Å^−3^. A constant integration timestep of 2 fs was utilized for every simulation in tandem with the SHAKE algorithm for the constraint of hydrogen atoms. To obtain a disordered membrane bilayer, we did minimizations of 1000 steps followed by equilibrations lasting 0.5 ns with all atoms except the lipid tails constrained (*k =* 1 kcal/mol/Å^2^) when indicated (Table 1). These simulations were followed by another minimization of 1000 steps and by an equilibration for another 0.5 ns with protein backbone atoms constrained (*k =* 1 kcal/mol/Å^2^). All simulations were performed at 310 K and 1 atmosphere of pressure by using the Langevin thermostat and the hybrid Nosé-Hoover piston method for pressure control. A Langevin damping coefficient of 0.1 ps^−1^ was chosen for the thermostat while a piston period of 200 fs and damping timescale of 50 fs was used for pressure control. Atomic coordinates were saved every 2 ps in all production runs. A constant electric field applied to all atoms was used in simulations with a transmembrane potential^88^.

### Simulation analysis procedures and tools

Average water concentration maps were computed using the VMD VolMap Tool with a resolution of 1 Å^3^. Average electrostatic maps were computed using the PME electrostatic plugin and an Ewald factor of 0.25 with a grid spacing > 1 Å^3^. Cell dimensions for PME calculations were set as their average for the entire simulation. Both volumetric density maps and electrostatic potential maps were computed every 50 ps with systems aligned using the protein backbone and starting coordinates as reference. Ionic currents were obtained by counting the number of ions that sequentially traversed from bulk electrolyte, to a region between the two membrane planes, to bulk electrolyte on the other side of the membrane. Membrane plane positions were defined by the average z-coordinate of the phosphorus atoms of the lipid head groups for each leaflet. Residue distances were computed between Cα atoms every 50 ps unless otherwise stated. The HOLE program^136^ was utilized to calculate pore dimensions using coordinates from simulations taken every 1 ns. Lipid atom names were changed using VMD to be recognizable by HOLE. Estimation of TMC1 pore radii was difficult due to the curvaceous nature of the permeation pathway and lipid dynamics. Therefore, outputs of pore profiles obtained from HOLE were first visualized in VMD for curation. Any profile that showed permeation pathways that were clearly implausible were discarded prior to further analyses. Residues that lined the pore more often at a given position were identified using HOLE’s output. The radial distribution function *g*(*r*) of lipid tails with respect to C_α_ atoms was calculated with the corresponding VMD plugin over 40 Å with values compute every 0.1 Å. Periodic boundary conditions were taken into consideration and selection was updated every frame. Electrostatic potential and water concentration maps were plotted in MATLAB, all other data were plotted using xmgrace, and molecular images were generated using VMD^123^.

### Bulk electrolyte simulations

Control simulations of bulk electrolyte^87,137^ (0.150 M KCl electrolyte prepared in VMD using the solvate and autoionize plugins; 96,243 atoms, 100 Å^3^; Table S1) were performed in duplicate with identical parameters to those described above. The system was minimized for 1000 steps followed by 20 ns of equilibration. Two configurations, at 10 ns and at 20 ns, were taken from this equilibration for subsequent simulations with an applied voltage. Each configuration had a set of three simulations with a varying bias of −0.500 V, −0.250 V, and −0.125 V. We obtained estimates for the induced current and resistivity of the 0.150 M KCl solution by counting the number of ions that traversed the *z* axis through an imaginary box of dimensions *l_x_* = *l_y_* = 50 Å and *l_z_* = 20, 40, or 60 Å. Currents were found to be roughly independent of *l_z_*. Hence, all calculations for conductance of bulk electrolyte were carried out with *l_z_* = 20. The resistivity *ρ_i_* for system *i* was calculated using:

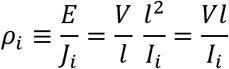

where *E* is the applied electric field computed as the ratio between the voltage and the length of simulation box in *z* direction (*V*/*l*), and *J_i_* is the current density (*I*_i_/*l*^2^). The average resistivity of a 0.150 M KCl solution under the simulated conditions was *ρ* = 0.278 Ohm m, which is equivalent to a conductivity of *σ* = 3.62 S m^−1^. The estimated experimental conductance of 0.150 M KCl at 308 K is 2.29 S m^−1^. Thus, currents from simulations are overestimated by ~ 63.3%. However, our simulations were carried out at 310 K, so we have used scaled conductance values (0.63*σ*) to report a lower boundary for the estimates.

## Supporting information

Supplementary Alignment

## ACKNOWLEDGEMENTS

We thank Nurunisa Akyuz, David P. Corey, Bechara Kachar, Jeffrey Holt, as well as all members of the Sotomayor laboratory for helpful discussions. This work was supported by the Ohio State University, by the National Institutes of Health (NIDCD R01 DC0155271), and by the Human Frontier Science Program (RGP0056/2018). Simulations were performed using the NCSA-Blue Waters (GLCPC), TACC-Stampede / PSC-Bridges (XSEDE MCB140226), OSC-Owens, and OSC-Pitzer (PAS1037 and PAA0217) supercomputers. CN was supported by an OSU/NIH molecular biophysics training grant (TG32GM118291). JML and JCS received Mayer’s summer undergraduate research fellowships. JML received an OSU undergraduate research scholarship. AB was supported by the Intramural Research Program NS002945 of the NINDS to Kenton J. Swartz.

## AUTHOR CONTRIBUTIONS

SW, JML, AB, and MS designed the research. SW, JML, CN, and JCS performed the simulations. SW and JML analyzed the data. SW, JML, and MS wrote the manuscript with feedback from all co-authors.

## SUPPLEMENTARY MATERIAL

### MOVIES

**Movie 1** Side view of the TMC1 channel pore at 0 V (monomer B). A potassium ion (pink) is seen exploring the pore without fully permeating from one membrane side to the other during an equilibrium simulation (Sim1a and Sim1b; 60 ns). Protein is shown in purple, phosphorus atoms from lipid head groups are shown as light-yellow spheres. Water molecules, lipid tails, and protein side chains are omitted for clarity.

**Movie 2** Side view of the TMC1 channel pore at −0.500 V (monomer A). Potassium (pink) and chloride ions (green) were observed to fully permeate from one membrane side to the other (Sim1c; 80 ns). System shown as in Movie 1.

**Fig. S1.**
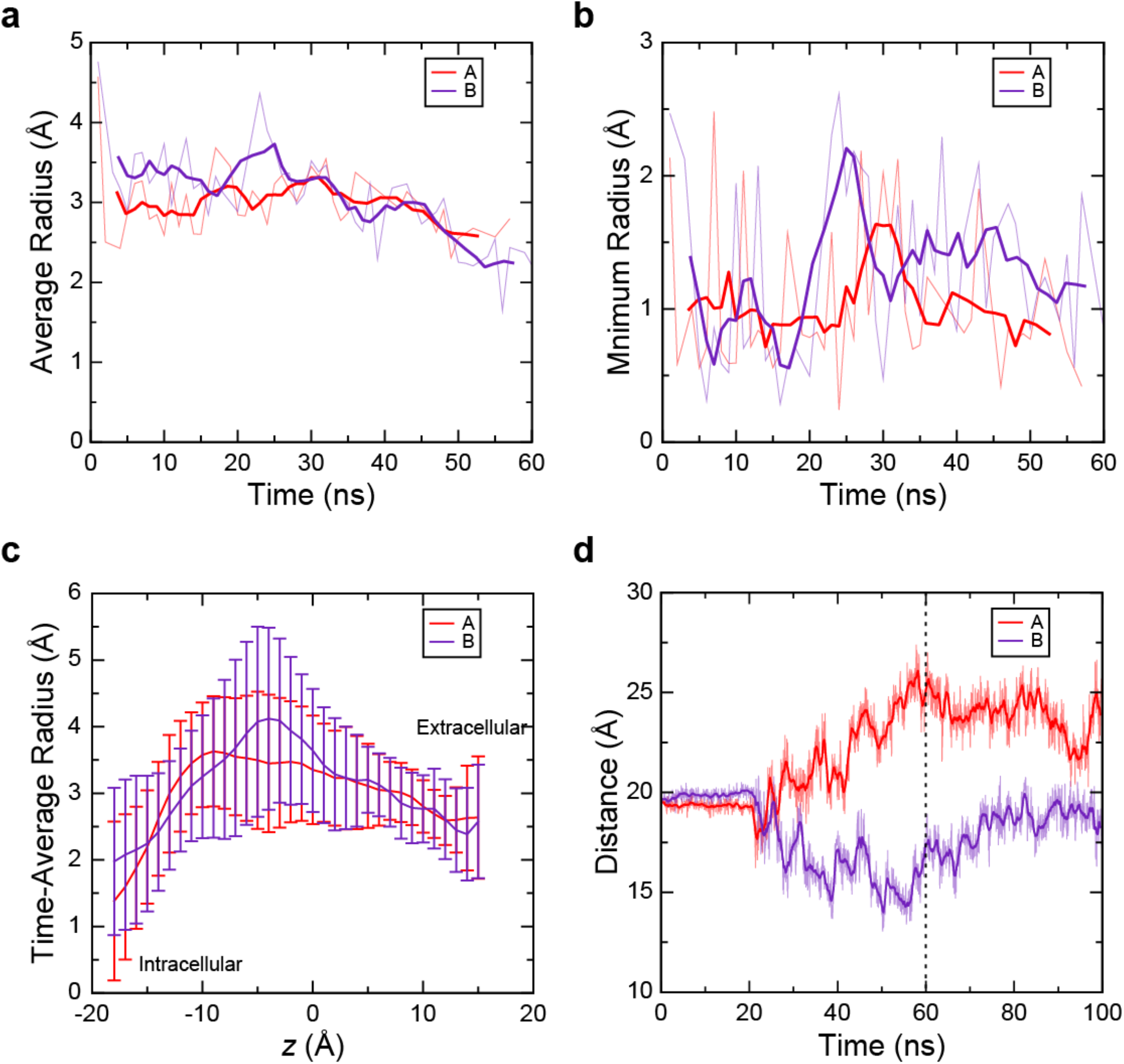
Pore properties during equilibration of full-length TMC1 model 1. **a** Average radius across the pore axis (*z*) for each monomer as a function of time during Sim1a (21 ns) and Sim1b (first 39 ns). Transparent lines are curated raw data and solid lines are their 5-ns running averages. **b** Minimum radii of each monomer pore as a function of time during the same 60 ns of simulation as in **a**. **c** Time-average radii as a function of position (*z*) along the pore axis computed over 60 ns of Sim1a and Sim1b. Value of *z* increases from intra- to extra-cellular sides. Error bars are standard deviation. **d** Separation between C_α_ atoms of Met^418^ in α4 and Thr^535^ in α6 (Sim1a and Sim1b). Transparent lines indicate raw data recorded every 50 ps, solid lines are 1 ns running averages. Vertical dashed line marks starting point for simulations with transmembrane potential (Sim1c and Sim1d).

**Fig. S2.**
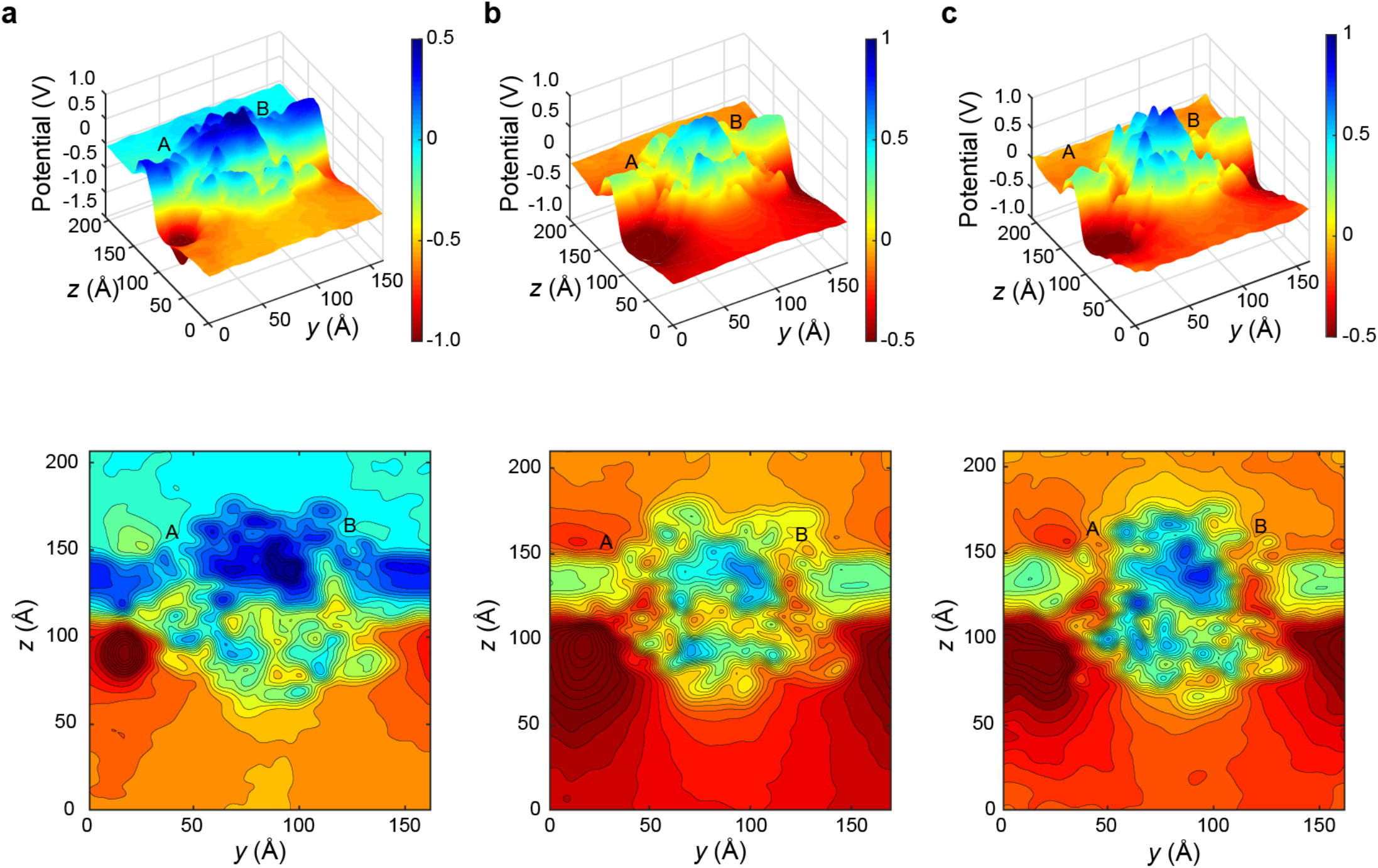
Electrostatic potential surface for the full-length TMC1 system. **a-c** Cross-section slice through TMC1 transmembrane pores showing the averaged electrostatic potential surface (top) and its two-dimensional contour plot for full length TMC1 **a** at −0.500 V (80 ns; Sim1c), **b** at −0.250 V (250 ns; Sim1d), and **c** at −0.125 V (100 ns; Sim1f).

**Fig. S3.**
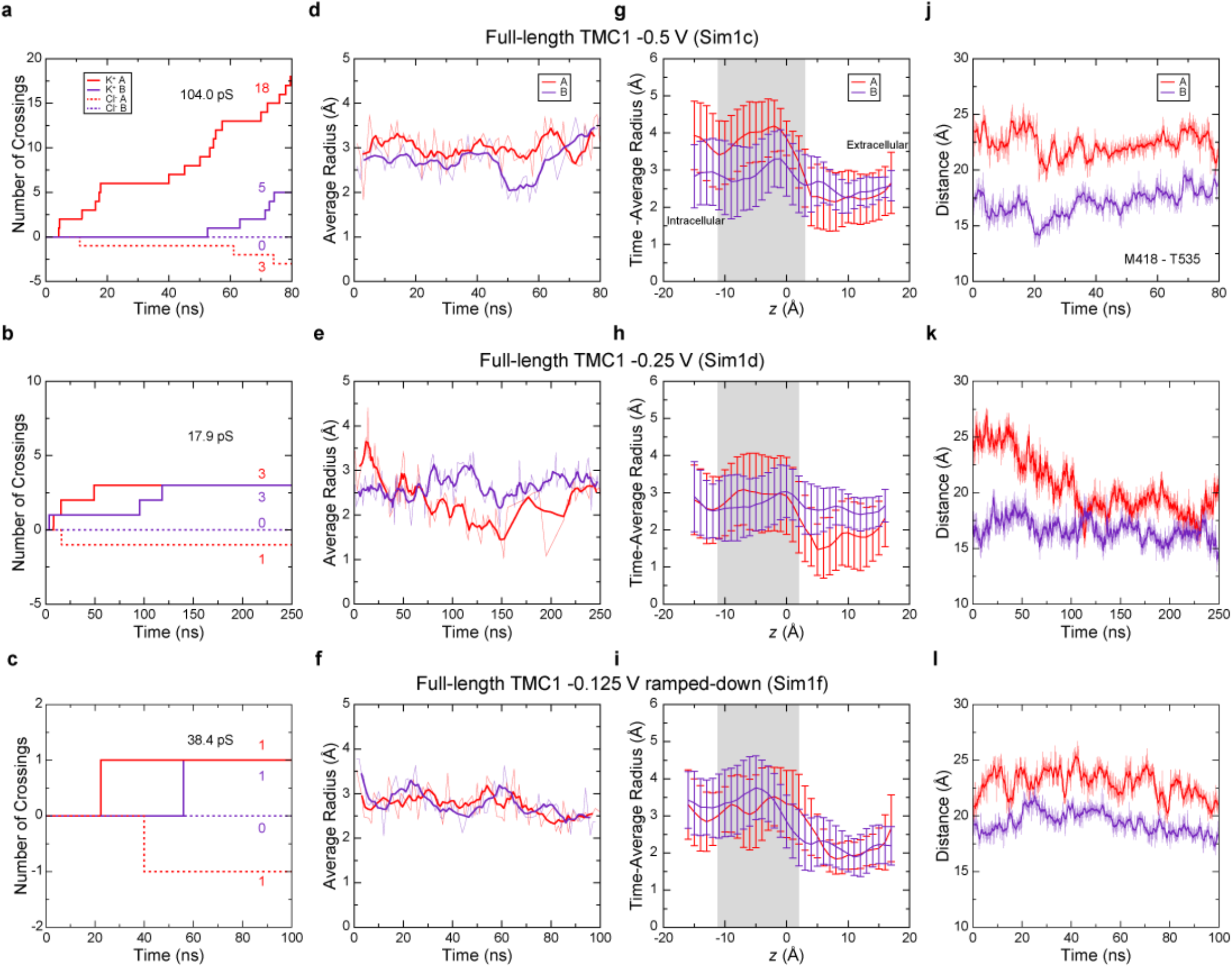
Ion conduction and pore properties of TMC1. **a-c** Number of ion crossings as a function of time for full-length TMC1 simulated at **a**-0.500 V (Sim1c), **b**-0.250 V (Sim1d), and **c**-0.125 V (Sim1f). Solid lines are for potassium ions going through monomers A (red) and B (indigo). Dashed lines are for chloride ions going through monomers A (red) and B (indigo). **d-f** Average radius across the pore axis (*z*) for each monomer as a function of time for TMC1 simulated at **d**-0.500 V (Sim1c), **e** at −0.250 V (Sim1d), and **f** at −0.125 V (Sim1f). Transparent lines show raw data after curation (see Methods) while solid lines indicate their 5-ns running averages. **g-i** Time-averaged radii as a function of position (*z*) along the pore axis for TMC1 simulated at **g**-0.500 V (Sim1c), **h**-0.250 V (Sim1d), and **i**-0.125 V (Sim1f). Value of *z* increases from intra- to extra-cellular sides. Error bars are standard deviation. **j-l** Separation between C_α_ atoms of Met^418^ in α4 and Thr^535^ in α6 as a function of time for TMC1 simulated at **j**-0.500 V (Sim1c), **k** at −0.250 V (Sim1d), and **l**-0.125 V (Sim1f). Transparent lines indicate raw data recorded every 50 ps, solid lines are 1 ns running averages.

**Fig. S4.**
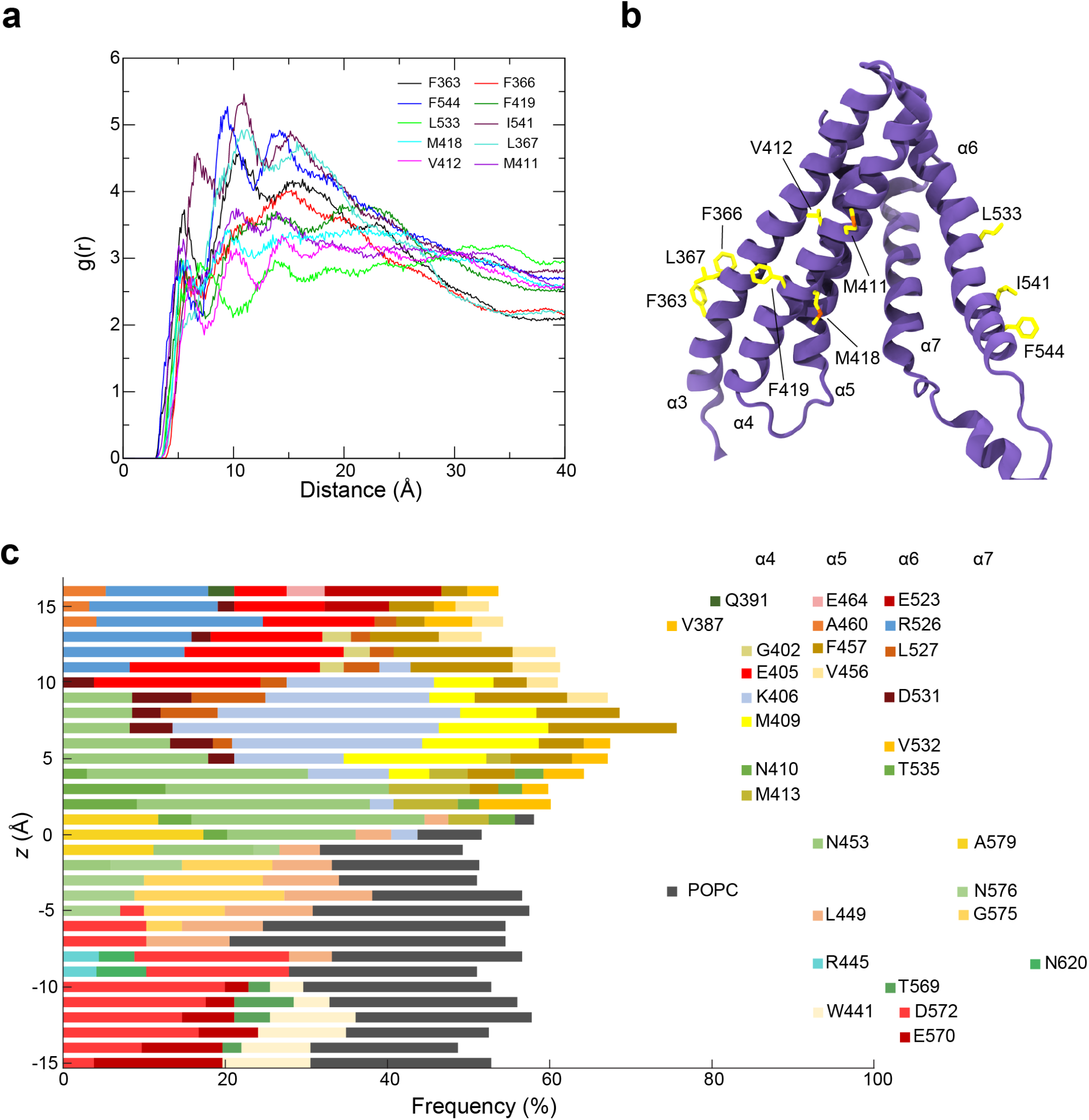
Membrane-interacting and pore-lining residues. **a** Radial distribution function *g*(*r*) for lipid tails as a function of distance from C_α_ atoms of key hydrophobic residues in monomer A during a simulation of TMC1 at −0.500 V (Sim1c). Top 10 residues were selected after computing *g*(*r*) for hydrophobic residues in helices α3 to α7 during Sim1c and computing the highest area under the curve between 0 to 6 Å. **b** TMC1 transmembrane helices α3 to α7 showing the location of residues listed in **a**. **c** Key residues that line the hydrophilic pore as a function of distance *z* along the pore axis during simulations of full-length TMC1 with transmembrane potential (Sim1d-Sim1f). To determine the frequency at which residues appeared to line the pore we analyzed the HOLE output and identified the three residues that lined the pore most often at each value of *z* along the axis in each of the simulations.

**Fig. S5.**
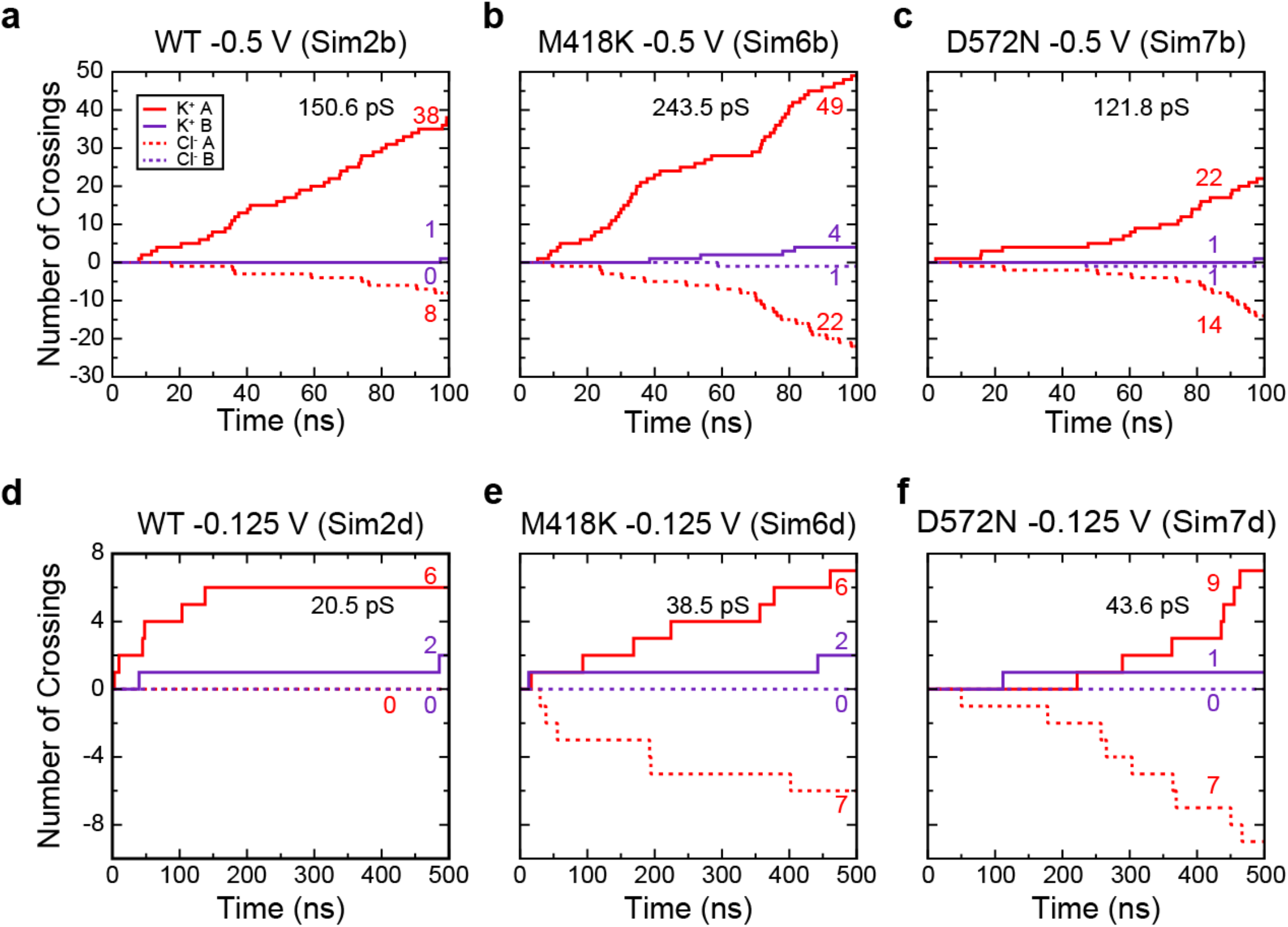
Ion conduction through a minimalistic truncated model of the TMC1 dimer. **a-c** Number of ion crossings through the pore of the wild type, M418K, and D572N truncated TMC1 systems at −0.500 V (Sim2b, Sim6b, and Sim7b). Solid and dashed lines denote potassium and chloride permeation events, respectively. **d-f** Ion crossing events through the pore of the wild type, M418K, and D572N truncated TMC1 systems at −0.125 V (Sim2d, Sim6d, and Sim7d). Solid and dashed lines denote potassium and chloride permeation events, respectively. Panels a,d are also shown in Fig. S7a,c.

**Fig. S6.**
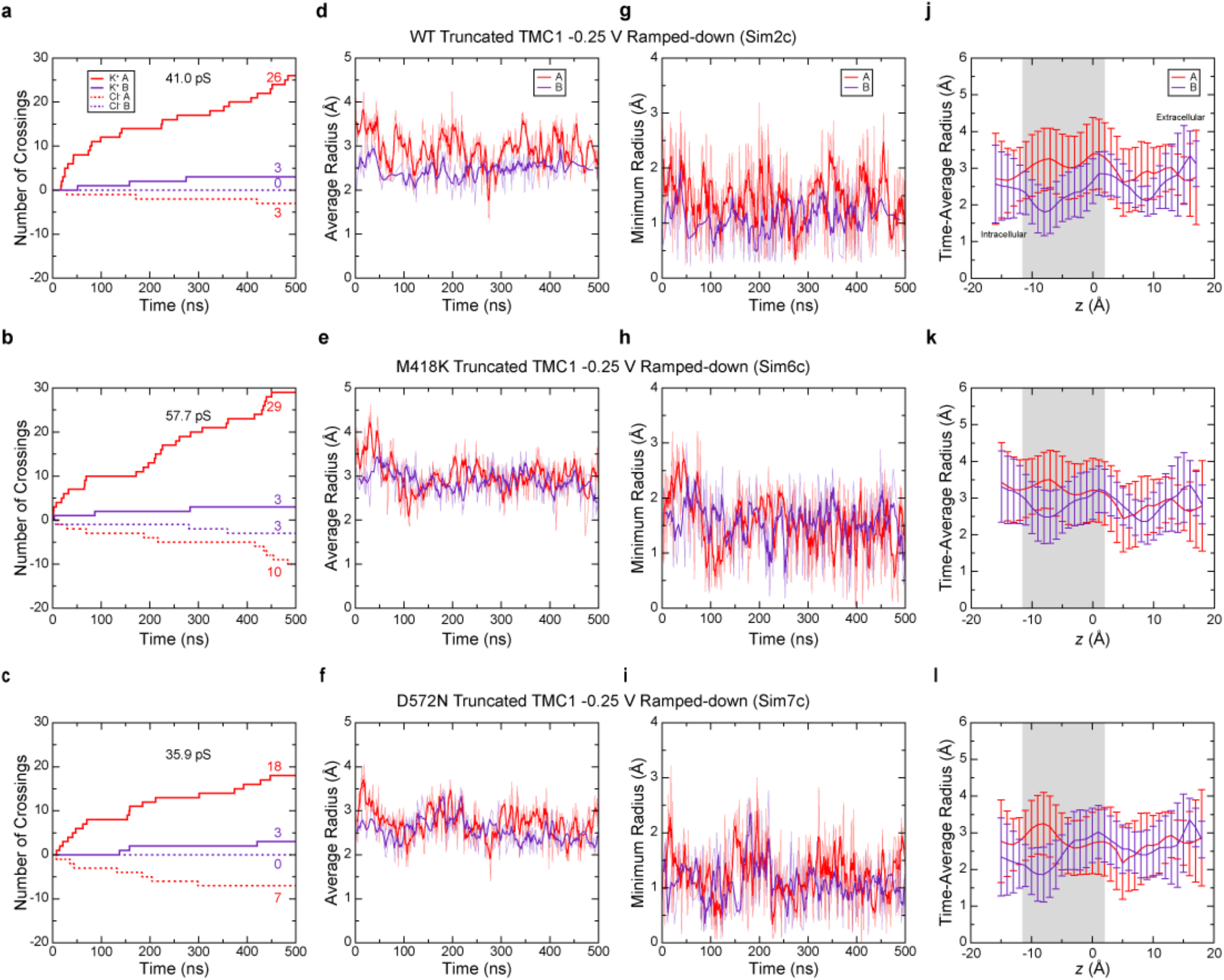
Ion conduction and pore properties of truncated TMC1 systems. **a-c** Number of ion crossings as a function of time for truncated TMC1 systems at −0.250 V including **a** wild-type protein (Sim2c; also shown in Fig. 3d), **b** M418K mutant (Sim6c), and **c** D572N mutant (Sim7c). Shown as in Fig. S3a-c. **d-f** Average radius across the pore axis (*z*) for each monomer as a function of time for truncated TMC1 systems at −0.250 V including **d** wild-type protein (Sim2c), **e** M418K mutant (Sim6c), and **f** D572N mutant (Sim7c). Shown as in Fig S3d-f. **g-i** Minimum radii of each monomer pore as a function of time for truncated TMC1 systems at −0.250 V including **g** wild-type protein (Sim2c), **h** M418K mutant (Sim6c), and **i** D572N mutant (Sim7c). **j-l** Time-averaged radii as a function of position (*z*) along the pore axis for truncated TMC1 systems at −0.250 V including **j** wild-type protein (Sim2c), **k** M418K mutant (Sim6c), and **l** D572N mutant (Sim7c). Shown as in Fig S3g-i. Panels a,d,g,j are also shown in Fig. S7b,e,h,k.

**Fig. S7.**
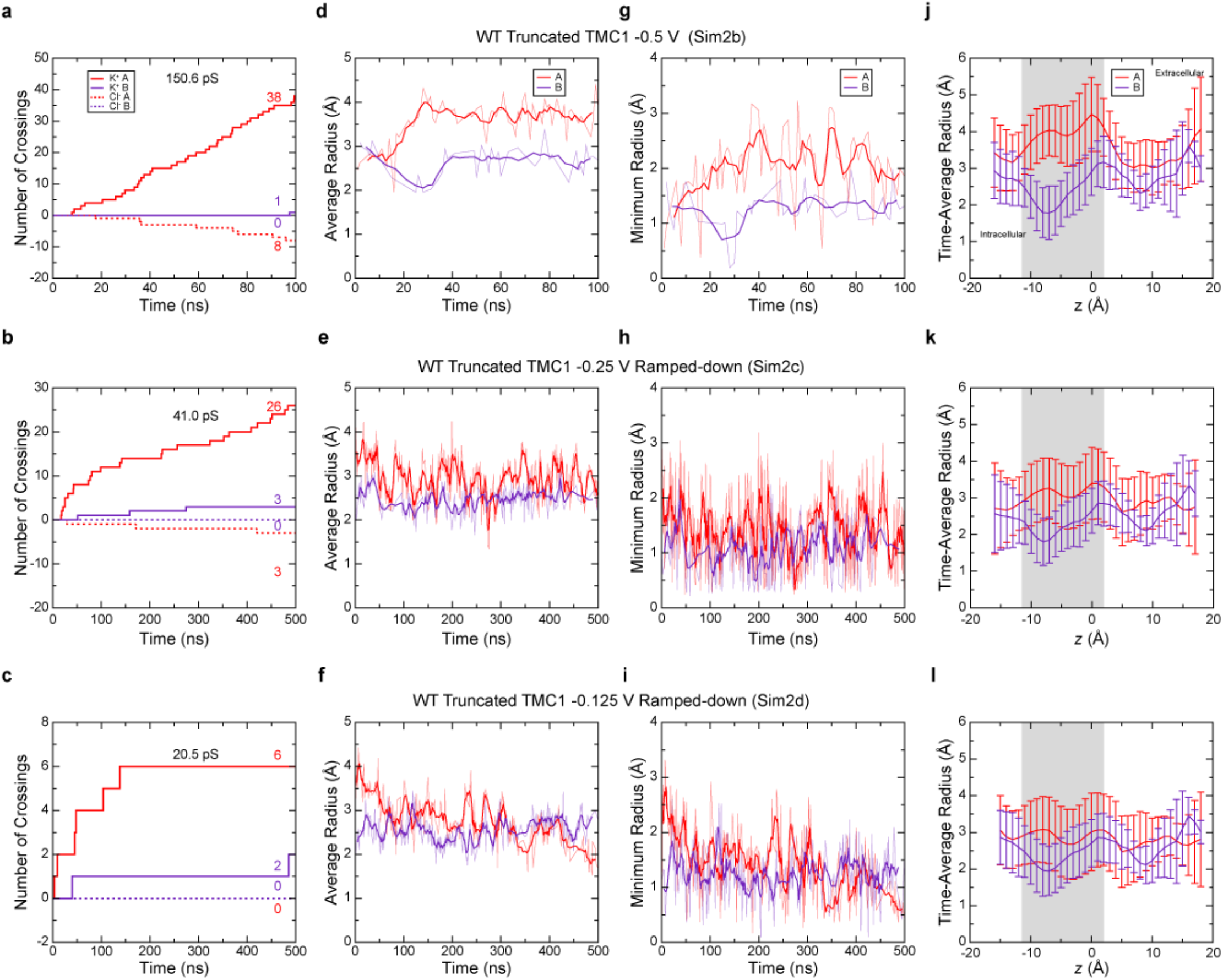
Ion conduction and pore properties of the wild-type truncated TMC1 system. **a-c** Number of ion crossings as a function of time for a wild-type truncated TMC1 system at **a**-0.500 V (Sim2b), **b**-0.250 V (Sim2c), and **c**-0.125 V (Sim2d). Shown as in Fig. S3a-c. **d-f** Average radius across the pore axis (*z*) for each monomer as a function of time for a wild-type truncated TMC1 system at **d**-0.500 V (Sim2b), **e**-0.250 V (Sim2c), and **f**-0.125 V (Sim2d). Shown as in Fig S3 d-f. **g-i** Minimum radii of each monomer pore as a function of time for a wild-type truncated TMC1 system at **g**-0.500 V (Sim2b), **h**-0.250 V (Sim2c), and **i**-0.125 V (Sim2d). **j-l** Time-averaged radii as a function of position (*z*) along the pore axis for a wild-type truncated TMC1 system at **j**-0.500 V (Sim2b), **k**-0.250 V (Sim2c), and **l**-0.125 V (Sim2d). Shown as in Fig S3g-i. Panels b,e,h,k are also shown in Fig. S6a,d,g,j. Panels a,c are also shown in Fig. S5a,d.

**Fig. S8.**
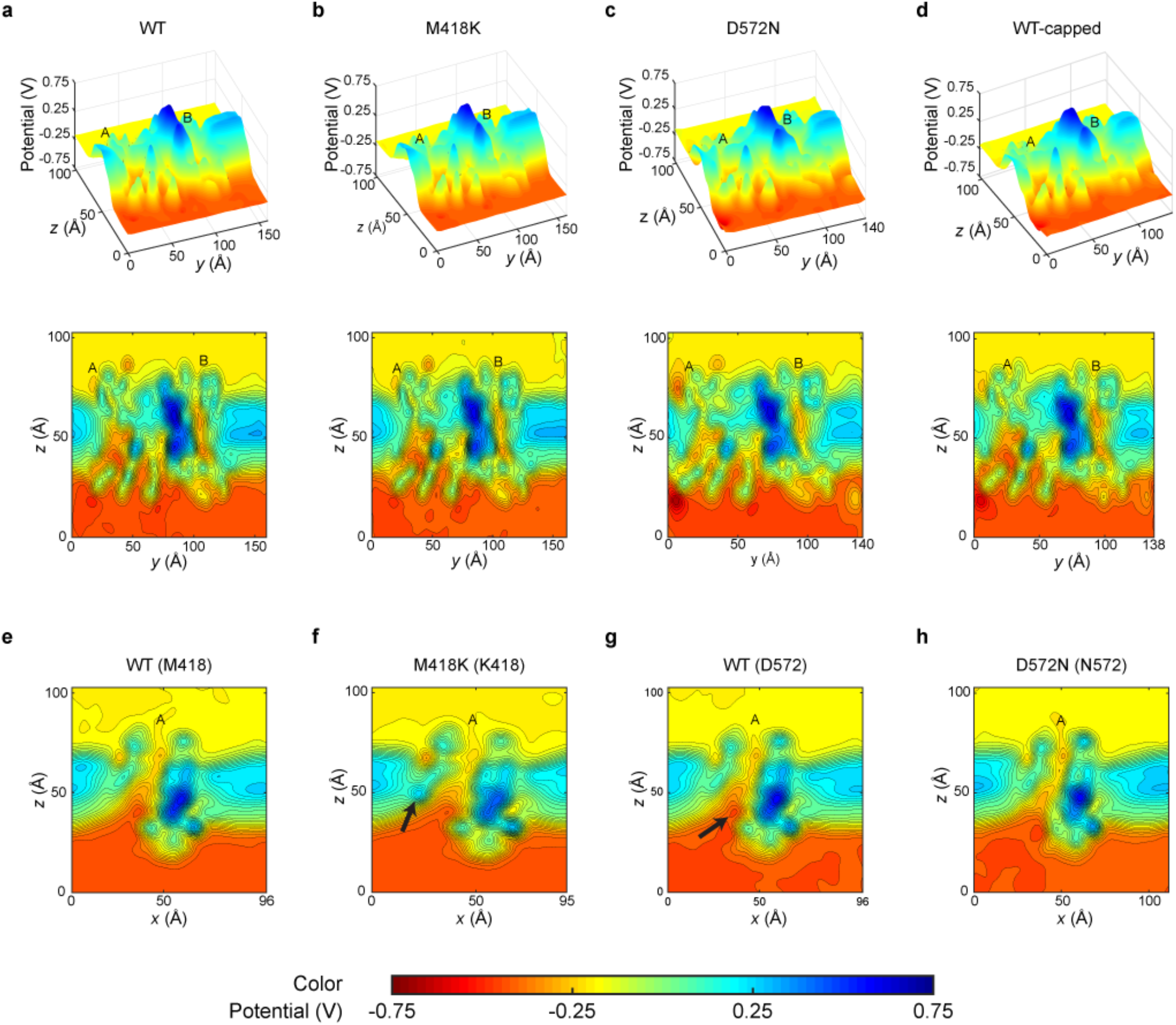
Electrostatic potential surface for truncated TMC1 systems. **a-d** Cross-section slice through TMC1 transmembrane pores showing the averaged electrostatic potential surface (top) and a its two-dimensional contour plot at - 0.250 V for the **a** wild-type protein (Sim2c), **b** M418K mutant (Sim6c), **c** D572N (Sim7c), and **d** wild-type capped protein (Sim8c). **e-h** Two-dimensional contour plot of a monomer A cross-section slices of the averaged electrostatic potential at - 0.250 V for **e** wild-type protein (plane passing through M418; Sim2c), **f** M418K mutant (plane passing through K418; Sim6c), **g** wild-type protein (plane passing through D572; Sim2c), and **h** D572N mutant (plane passing through N572; Sim7c). Black arrows indicate changes in electrostatic potential near the mutated residues.

**Table S1.** Control simulations of KCl bulk electrolyte.

| Label | Length (ns) | Voltage | $K^+_{+z}$ | $K^+_{-z}$ | $Cl^-_{+z}$ | $Cl^-_{-z}$ | I pA | C (pS) |
| --- | --- | --- | --- | --- | --- | --- | --- | --- |
| Sim9a | 10 | 0 | - | - | - | - | - | - |
| Sim9b | 10 | 0 | - | - | - | - | - | - |
| Sim9c | 10 | -0.500 | 11 | 589 | 558 | 17 | 17.93 | 35.85 |
| Sim9d | 10 | -0.500 | 11 | 569 | 599 | 12 | 18.34 | 36.69 |
| Sim9e | 10 | -0.250 | 58 | 335 | 334 | 50 | 8.99 | 35.95 |
| Sim9f | 10 | -0.250 | 58 | 346 | 329 | 56 | 8.99 | 35.95 |
| Sim9g | 10 | -0.125 | 90 | 243 | 240 | 92 | 4.82 | 38.58 |
| Sim9h | 10 | -0.125 | 97 | 231 | 242 | 110 | 4.26 | 34.09 |

## Notes

### Competing Interest Statement

The authors have declared no competing interest.

