## Supplementary Alignment for "*In Silico* Electrophysiology of Inner-Ear Mechanotransduction Channel TMC1 Models"

```
Hs         ------------------------------------------------------------------------
Mm         ------------------------------------------------------------------------
Pt         ------------------------------------------------------------------------
Ss         ------------------------------------------------------------------------
Bt         ------------------------------------------------------------------------
Rn         ------------------------------------------------------------------------
Clf        ------------------------------------------------------------------------
Fc      1  ------------------------MGWDYVPLTPAVA--------QTP-TLIKRPTYLQQRATLSISPLSLE   39
Gg      1  -----------------------MAVLELSLLMHVKLFSKDLALINDN-TVL---------RFLG--KNGSD   37
Dr      1  MLVKSELFMHATHHRQALFSQAMVYCCDIIIIIIIIIIIITHTHGQPHARLLRR-RMPRHKLIAS--ESDVS   69
Am      1  -----------------------------------------------M-TVH---------RFKC-----FK   10
Cm      1  -------------------------MYDYVSILEIVFLSSGMSPGGDMSQPS---------SFQP--GVGIE   36

Hs         ------------------------------------------------------------------------
Mm         ------------------------------------------------------------------------
Pt         ------------------------------------------------------------------------
Ss         ------------------------------------------------------------------------
Bt      1  -------------------------------------------------MIRH----------FIFVLDSSG   13
Rn         ------------------------------------------------------------------------
Clf        ------------------------------------------------------------------------
Fc     40  IGAPLGPVE-S------VDPPNAPASVPVAPA-------ERQRGDSRWCQLHRWVKGGEPPTCTAVAIQPNT   97
Gg     38  VNNEKDSAAENQENGRGADKERK----RAKP--------RTRRGHARGRARHGGEEDGDDE------DGTRR   91
Dr     70  IEVDEGKDKESCVYY--VEVEENCERGKIKQASRDGKRRRERNGETRRKASEKRTNEGESKK----AEKKHE  135
Am     11  VEVEEGSAAESEEND------------------------------KTGENKTYGDDEGDQK------PDP--   44
Cm     37  VDNEEGSAAESEDDDSKIDEEESEKEKKVTA--------KTRRGNARGKARK----DSEEE------EETRT   90

Hs      1  ---------------------MSPKKVQIKVE--EKEDETEESS---------------SEEEEEVEDKLPR   34
Mm      1  -------------------------MLQIQVE--EKEEDTEESS---------------SE---EEEDKLPR   27
Pt      1  ---------------------MPPKKVQIKVE--EKEDETEESS---------------SEEEEEEEDKLPR   34
Ss      1  ---------------------MPPKKVQIQVE--EKEDDTEESS---------------SG--EEEENKLPR   32
Bt     14  GFVWEGLLSLVLQHYFEFQEMQISIDMQIQVE--EKEDDTEESS---------------SEE-EEEENKLPR   67
Rn         ------------------------------------------------------------------------
Clf        ------------------------------------------------------------------------
Fc     98  DTIWMET--GMNHAWLPIGWTVSSLAVQIQVE--EKEDDTEESS---------------SEEEEEEEDKLPR  150
Gg     92  AGQRRQR------NR----------RGRGRAGGNEGSDSEGDPTKARKNQANQKSKE-DD--EQEEDEKGGK  144
Dr    136  KGHRTARKAGEKHGK----------RQRRKNAGEE--DAEDKSSKEKKNMKNEKNKTLKLEEEKEKDVRKKK  195
Am     45  -VHRVPR------KR----------PQRIRVG--LESDSDEEVSKSRKNRANQKSKE-EGGEDGEDEEKGGK   96
Cm     91  AGQHRQR------NR----------RAQRKQAGNEDSDSDGESCKNKKNRENQISKQQEPNEDGEEDNKGGR  146

Hs     35  RESLRPKRKR------------TRDVINEDDPEPEPEDEETRKA-REKERRRRLKRGAEE-EEIDEEELERL   92
Mm     28  RESLRPKRKR------------TRDVINEDDPEPEPEDEETRKA-REKERRRRLRRGAEEEEEIDEEELERL   86
Pt     35  RESLRPKRKR------------TRDVINEDDPEPEPEDEETRKA-REKERRRRLKRGAE-------------   80
Ss     33  RESLRPKRKR------------TRDVINEDDPEPEPEDEETRKA-REKERRRRLRRGAEEEEEIDEEELERL   91
Bt     68  RQSLRPKRKR------------TRDVINEDDPEPEPEDEETRKA-REKERRRRLKRGAEEEEEIDAEELERL  126
Rn         ------------------------------------------------------------------------
Clf        ------------------------------------------------------------------------
Fc    151  RESLRPKRKR------------KRDVIHEDDPEPEPEDEETRKA-REKERRRRLKRGAEEEEEIDEEELERL  209
Gg    145  SGKGKNEKKQDKK-ADKKGTKEKANKKKK----------DKKKSGSGSDSDSE-------EEPITEEELAKL  198
Dr    196  RKHVKNEEDETNHEKTKQHLKEEKRRKKRKKPETTSESESKSESESASESESKNSPAVGVLGSLTPEELENL  267
Am     97  KGRGKNDK----K-NDKKGKKEKANKKKK----------DKKKSGSGSDSDSE-------EEPITEEELAKL  146
Cm    147  KGRGKNETKQEKK-PDEKGKKEKDNKKKD----------KKKKGSSSSDSDSE-------EEPITEEELAKL  200
```

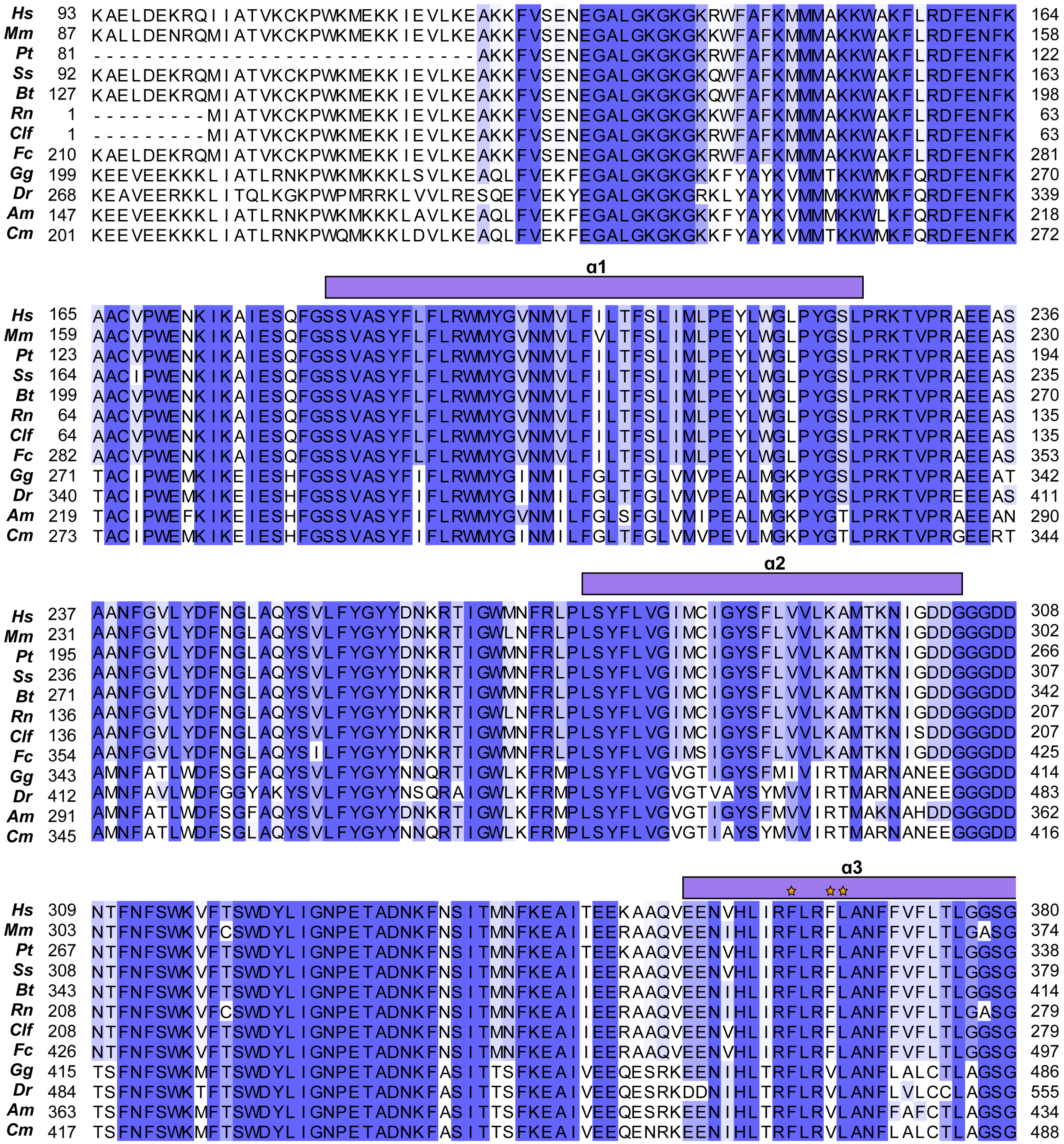
Hs 93 KAELDEKRQIIATVKCKPWKMEKKIEVLKEAKKFVSENEGALGKGKGKRWFAFKMMMAKKWAKFLRDFENFK 164
Mm 87 KALLDENRQMIATVKCKPWKMEKKIEVLKEAKKFVSENEGALGKGKGKKWFAFKMMMAKKWAKFLRDFENFK 158
Pt 81 ------------------------------AKKFVSENEGALGKGKGKRWFAFKMMMAKKWAKFLRDFENFK 122
Ss 92 KAELDEKRQMIATVKCKPWKMEKKIEVLKEAKKFVSENEGALGKGKGKQWFAFKMMMAKKWAKFLRDFENFK 163
Bt 127 KAELDEKRQMIATVKCKPWKMEKKIEVLKEAKKFVSENEGALGKGKGKQWFAFKMMMAKKWAKFLRDFENFK 198
Rn 1 ---------MIATVKCKPWKMEKKIEVLKEAKKFVSENEGALGKGKGKKWFAFKMMMAKKWAKFLRDFENFK 63
Clf 1 ---------MIATVKCKPWKMEKKIEVLKEAKKFVSENEGALGKGKGKRWFAFKMMMAKKWAKFLRDFENFK 63
Fc 210 KAELDEKRQMIATVKCKPWKMEKKIEVLKEAKKFVSENEGALGKGKGKRWFAFKMMMAKKWAKFLRDFENFK 281
Gg 199 KEEVEEKKKLIATLRNKPWKMKKKLSVLKEAQLFVEKFEGALGKGKGKKFYAYKVMMTKKWMKFQRDFENFK 270
Dr 268 KEAVEERKKLITQLKGKPWPMRRKLVVLRESQEFVEKYEGALGKGKGRKLYAYKVMMMKKWMKFQRDFENFK 339
Am 147 KEEVEEKKKLIATLRNKPWKMKKKLAVLKEAQLFVEKFEGALGKGKGKKFYAYKVMMMKKWLKFQRDFENFK 218
Cm 201 KEEVEEKKKLIATLRNKPWQMKKKLDVLKEAQLFVEKFEGALGKGKGKKFYAYKVMMTKKWMKFQRDFENFK 272
α1
Hs 165 AACVPWENKIKAIESQFGSSVASYFLFLRWMYGVNMVLFILTFSLIMLPEYLWGLPYGSLPRKTVPRAEEAS 236
Mm 159 AACVPWENKIKAIESQFGSSVASYFLFLRWMYGVNMVLFVLTFSLIMLPEYLWGLPYGSLPRKTVPRAEEAS 230
Pt 123 AACVPWENKIKAIESQFGSSVASYFLFLRWMYGVNMVLFILTFSLIMLPEYLWGLPYGSLPRKTVPRAEEAS 194
Ss 164 AACIPWENKIKAIESQFGSSVASYFLFLRWMYGVNMVLFILTFSLIMLPEYLWGLPYGSLPRKTVPRAEEAS 235
Bt 199 AACVPWENKIKAIESQFGSSVASYFLFLRWMYGVNMVLFILTFSLIMLPEYLWGLPYGSLPRKTVPRAEEAS 270
Rn 64 AACVPWENKIKAIESQFGSSVASYFLFLRWMYGVNMVLFILTFSLIMLPEYLWGLPYGSLPRKTVPRAEEAS 135
Clf 64 AACVPWENKIKAIESQFGSSVASYFLFLRWMYGVNMVLFILTFSLIMLPEYLWGLPYGSLPRKTVPRAEEAS 135
Fc 282 AACVPWENKIKAIESQFGSSVASYFLFLRWMYGVNMVLFILTFSLIMLPEYLWGLPYGSLPRKTVPRAEEAS 353
Gg 271 TACIPWEMKIKEIESHFGSSVASYFIFLRWMYGINMILFGLTFGLVMVPEALMGKPYGSLPRKTVPRAEEAT 342
Dr 340 TACIPWEMKIKEIESHFGSSVASYFIFLRWMYGINMILFGLTFGLVMVPEALMGKPYGSLPRKTVPREEEAS 411
Am 219 TACIPWEFKIKEIESHFGSSVASYFIFLRWMYGVNMILFGLSFGLVMIPEALMGKPYGTLPRKTVPRAEEAN 290
Cm 273 TACIPWEMKIKEIESHFGSSVASYFIFLRWMYGINMILFGLTFGLVMVPEVLMGKPYGTLPRKTVPRGEERT 344
α2
Hs 237 AANFGVLYDFNGLAQYSVLFYGYYDNKRTIGWMNFRLPLSYFLVGIMCIGYSFLVVLKAMTKNIGDDGGGDD 308
Mm 231 AANFGVLYDFNGLAQYSVLFYGYYDNKRTIGWLNFRLPLSYFLVGIMCIGYSFLVVLKAMTKNIGDDGGGDD 302
Pt 195 AANFGVLYDFNGLAQYSVLFYGYYDNKRTIGWMNFRLPLSYFLVGIMCIGYSFLVVLKAMTKNIGDDGGGDD 266
Ss 236 AANFGVLYDFNGLAQYSVLFYGYYDNKRTIGWMNFRLPLSYFLVGIMCIGYSFLVVLKAMTKNIGDDGGGDD 307
Bt 271 AANFGVLYDFNGLAQYSVLFYGYYDNKRTIGWMNFRLPLSYFLVGIMCIGYSFLVVLKAMTKNIGDDGGGDD 342
Rn 136 AANFGVLYDFNGLAQYSVLFYGYYDNKRTIGWLNFRLPLSYFLVGIMCIGYSFLVVLKAMTKNIGDDGGGDD 207
Clf 136 AANFGVLYDFNGLAQYSVLFYGYYDNKRTIGWMNFRLPLSYFLVGIMCIGYSFLVVLKAMTKNISDDGGGDD 207
Fc 354 AANFGVLYDFNGLAQYSILFYGYYDNKRTIGWMNFRLPLSYFLVGIMSIGYSFLVVLKAMTKNIGDDGGGDD 425
Gg 343 AMNFATLWDFSGFAQYSVLFYGYYNNQRTIGWLKFRMPLSYFLVGVGTIGYSFMIVIRTMARNANEEGGGDD 414
Dr 412 AMNFAVLWDFGGYAKYSVLFYGYYNSQRAIGWLKFRMPLSYFLVGVGTVAYSYMVVIRTMARNANEEGGGDD 483
Am 291 AMNFATLWDFSGFAQYSVLFYGYYDNKRTIGWLKFRMPLSYFLVGVGTIGYSFMVVIRTMAKNAHDDGGGDD 362
Cm 345 AMNFATLWDFSGLAQYSVLFYGYYNNQRTIGWLKFRMPLSYFLVGVGTIAYSYMVVIRTMARNANEEGGGDD 416
α3
Hs 309 NTFNFSWKVFTSWDYLIGNPETADNKFNSITMNFKEAITEEKAAQVEENVHLIRFLRFLANFFVFLTLGGSG 380
Mm 303 NTFNFSWKVFCSWDYLIGNPETADNKFNSITMNFKEAIIEERAAQVEENIHLIRFLRFLANFFVFLTLGASG 374
Pt 267 NTFNFSWKVFTSWDYLIGNPETADNKFNSITMNFKEAITEEKAAQVEENVHLIRFLRFLANFFVFLTLGGSG 338
Ss 308 NTFNFSWKVFTSWDYLIGNPETADNKFNSITMNFKEAIIEERAAQVEENVHLIRFLRFLANFFVFLTLGGSG 379
Bt 343 NTFNFSWKVFTSWDYLIGNPETADNKFNSITMNFKEAIIEERAAQVEENIHLIRFLRFLANFFVFLTLGGSG 414
Rn 208 NTFNFSWKVFCSWDYLIGNPETADNKFNSITMNFKEAIIEERAAQVEENIHLIRFLRFLANFFVFLTLGASG 279
Clf 208 NTFNFSWKVFTSWDYLIGNPETADNKFNSITMNFKEAIIEERAAQVEENVHLIRFLRFLANFFVFLTLGGSG 279
Fc 426 NTFNFSWKVFTSWDYLIGNPETADNKFNSITMNFKEAIIEERAAQVEENVHLIRFLRFLANFFVFLTLGGSG 497
Gg 415 TSFNFSWKMFTSWDYLIGNPETADNKFASITTSFKEAIVEEQESRKEENIHLTRFLRVLANFLALCTLAGSG 486
Dr 484 TSFNFSWKTFTSWDYLIGNPETADNKFASITTSFKEAIVEEQESRKDDNIHLTRFLRVLANFLVLCCLAGSG 555
Am 363 TSFNFSWKMFTSWDYLIGNPETADNKFASITTSFKEAIVEEQESRKEENIHLTRFLRVLANFFAFCTLAGSG 434
Cm 417 TSFNFSWKMFTSWDYLIGNPETADNKFASITTSFKEAIVEEQENRKEENIHLTRFLRVLANFLALCTLAGSG 488

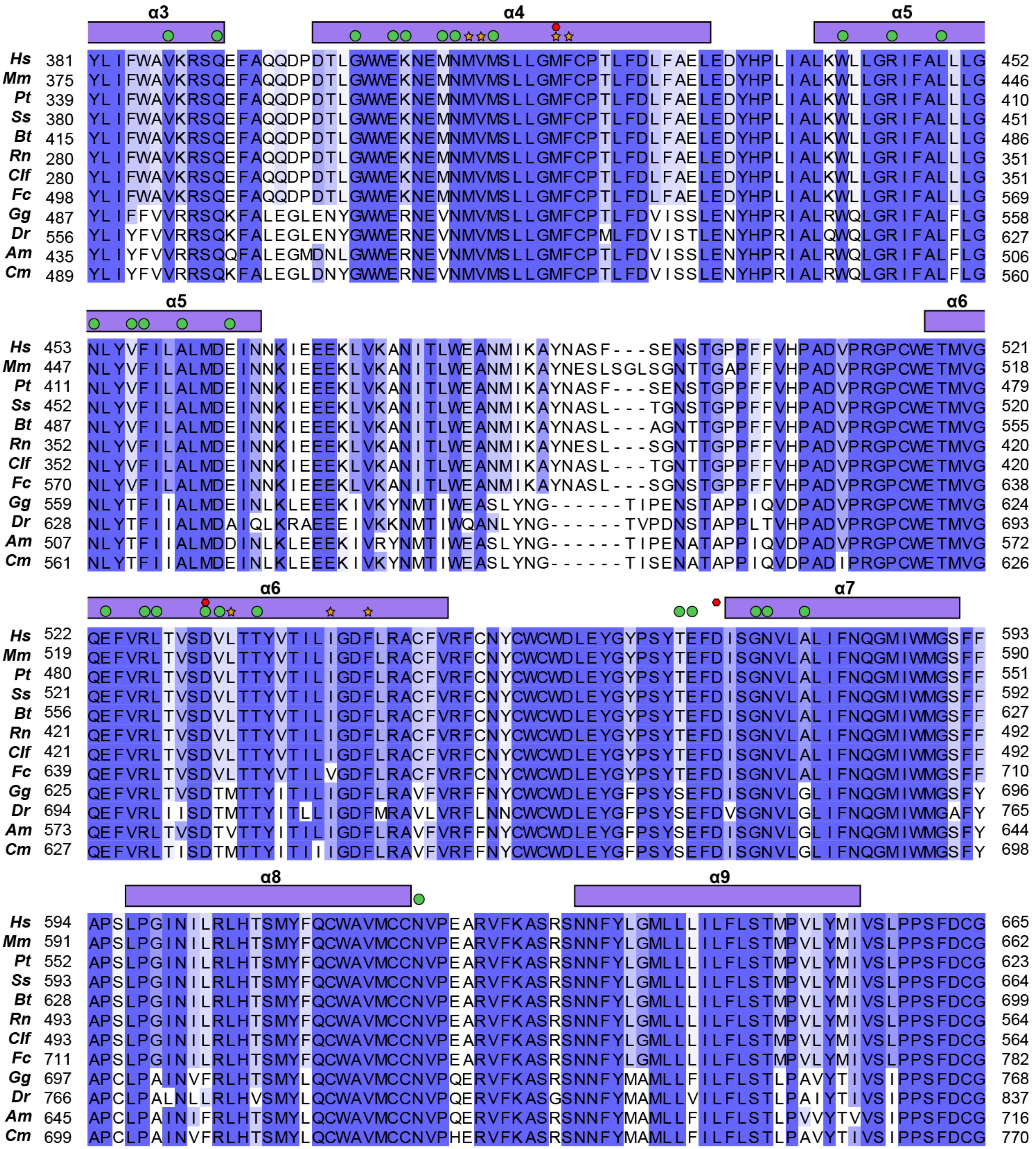
α3
α4
α5
Hs 381 YLIFWAVKRSQEFAQQDPDTLGWWEKNEMNMVMSLLGMFCPTLFDLFAELEDYHPLIALKWLLGRIFALLLG 452
Mm 375 YLIFWAVKRSQEFAQQDPDTLGWWEKNEMNMVMSLLGMFCPTLFDLFAELEDYHPLIALKWLLGRIFALLLG 446
Pt 339 YLIFWAVKRSQEFAQQDPDTLGWWEKNEMNMVMSLLGMFCPTLFDLFAELEDYHPLIALKWLLGRIFALLLG 410
Ss 380 YLIFWAVKRSQEFAQQDPDTLGWWEKNEMNMVMSLLGMFCPTLFDLFAELEDYHPLIALKWLLGRIFALLLG 451
Bt 415 YLIFWAVKRSQEFAQQDPDTLGWWEKNEMNMVMSLLGMFCPTLFDLFAELEDYHPLIALKWLLGRIFALLLG 486
Rn 280 YLIFWAVKRSQEFAQQDPDTLGWWEKNEMNMVMSLLGMFCPTLFDLFAELEDYHPLIALKWLLGRIFALLLG 351
Clf 280 YLIFWAVKRSQEFAQQDPDTLGWWEKNEMNMVMSLLGMFCPTLFDLFAELEDYHPLIALKWLLGRIFALLLG 351
Fc 498 YLIFWAVKRSQEFAQQDPDTLGWWEKNEMNMVMSLLGMFCPTLFDLFAELEDYHPLIALKWLLGRIFALLLG 569
Gg 487 YLIFFVVRRSQKFALEGLENYGWWERNEVNMVMSLLGMFCPTLFDVISSLENYHPRIALRWQLGRIFALFLG 558
Dr 556 YLIYFVVRRSQKFALEGLENYGWWERNEVNMVMSLLGMFCPMLFDVISTLENYHPRIALQWQLGRIFALFLG 627
Am 435 YLIYFVVRRSQQFALEGMDNLGWWERNEVNMVMSLLGMFCPTLFDVISSLENYHPRIALRWQLGRIFALFLG 506
Cm 489 YLIYFVVRRSQKFALEGLDNYGWWERNEVNMVMSLLGMFCPTLFDVISSLENYHPRIALRWQLGRIFALFLG 560
α5
α6
Hs 453 NLYVFILALMDEINNKIEEEKLVKANITLWEANMIKAYNASF---SENSTGPPFFVHPADVPRGPCWETMVG 521
Mm 447 NLYVFILALMDEINNKIEEEKLVKANITLWEANMIKAYNESLSGLSGNTTGAPFFVHPADVPRGPCWETMVG 518
Pt 411 NLYVFILALMDEINNKIEEEKLVKANITLWEANMIKAYNASF---SENSTGPPFFVHPADVPRGPCWETMVG 479
Ss 452 NLYVFILALMDEINNKIEEEKLVKANITLWEANMIKAYNASL---TGNSTGPPFFVHPADVPRGPCWETMVG 520
Bt 487 NLYVFILALMDEINNKIEEEKLVKANITLWEANMIKAYNASL---AGNTTGPPFFVHPADVPRGPCWETMVG 555
Rn 352 NLYVFILALMDEINNKIEEEKLVKANITLWEANMIKAYNESL---SGNTTGPPFFVHPADVPRGPCWETMVG 420
Clf 352 NLYVFILALMDEINNKIEEEKLVKANITLWEANMIKAYNASL---TGNTTGPPFFVHPADVPRGPCWETMVG 420
Fc 570 NLYVFILALMDEINNKIEEEKLVKANITLWEANMIKAYNASL---SGNSTGPPFFVHPADVPRGPCWETMVG 638
Gg 559 NLYTFIIALMDEINLKLEEEKIVKYNMTIWEASLYNG------TIPENSTAPPIQVDPADVPRGPCWETMVG 624
Dr 628 NLYTFIIALMDAIQLKRAEEEIVKKNMTIWQANLYNG------TVPDNSTAPPLTVHPADVPRGPCWETMVG 693
Am 507 NLYTFIIALMDDINLKLEEEKIVRYNMTIWEASLYNG------TIPENATAPPIQVDPADVPRGPCWETMVG 572
Cm 561 NLYTFIIALMDEINLKLEEEKIVKYNMTIWEASLYNG------TISENATAPPIQVDPADIPRGPCWETMVG 626
α6
α7
Hs 522 QEFVRLTVSDVLTTYVTILIGDFLRACFVRFCNYCWCWDLEYGYPSYTEFDISGNVLALIFNQGMIWMGSFF 593
Mm 519 QEFVRLTVSDVLTTYVTILIGDFLRACFVRFCNYCWCWDLEYGYPSYTEFDISGNVLALIFNQGMIWMGSFF 590
Pt 480 QEFVRLTVSDVLTTYVTILIGDFLRACFVRFCNYCWCWDLEYGYPSYTEFDISGNVLALIFNQGMIWMGSFF 551
Ss 521 QEFVRLTVSDVLTTYVTILIGDFLRACFVRFCNYCWCWDLEYGYPSYTEFDISGNVLALIFNQGMIWMGSFF 592
Bt 556 QEFVRLTVSDVLTTYVTILIGDFLRACFVRFCNYCWCWDLEYGYPSYTEFDISGNVLALIFNQGMIWMGSFF 627
Rn 421 QEFVRLTVSDVLTTYVTILIGDFLRACFVRFCNYCWCWDLEYGYPSYTEFDISGNVLALIFNQGMIWMGSFF 492
Clf 421 QEFVRLTVSDVLTTYVTILIGDFLRACFVRFCNYCWCWDLEYGYPSYTEFDISGNVLALIFNQGMIWMGSFF 492
Fc 639 QEFVRLTVSDVLTTYVTILVGDFLRACFVRFCNYCWCWDLEYGYPSYTEFDISGNVLALIFNQGMIWMGSFF 710
Gg 625 QEFVRLTVSDTMTTYITILIGDFLRAVFVRFFNYCWCWDLEYGFPSYSEFDISGNVLGLIFNQGMIWMGSFY 696
Dr 694 QEFVRLIISDTMTTYITLLIGDFMRAVLVRFLNNCWCWDLEYGFPSYSEFDVSGNVLGLIFNQGMIWMGAFY 765
Am 573 QEFVRLTVSDTVTTYITILIGDFLRAVFVRFFNYCWCWDLEYGFPSYSEFDISGNVLGLIFNQGMIWMGSFY 644
Cm 627 QEFVRLTISDTMTTYITIIIGDFLRAVFVRFFNYCWCWDLEYGFPSYSEFDISGNVLGLIFNQGMIWMGSFY 698
α8
α9
Hs 594 APSLPGINILRLHTSMYFQCWAVMCCNVPEARVFKASRSNNFYLGMLLLILFLSTMPVLYMIVSLPPSFDCG 665
Mm 591 APSLPGINILRLHTSMYFQCWAVMCCNVPEARVFKASRSNNFYLGMLLLILFLSTMPVLYMIVSLPPSFDCG 662
Pt 552 APSLPGINILRLHTSMYFQCWAVMCCNVPEARVFKASRSNNFYLGMLLLILFLSTMPVLYMIVSLPPSFDCG 623
Ss 593 APSLPGINILRLHTSMYFQCWAVMCCNVPEARVFKASRSNNFYLGMLLLILFLSTMPVLYMIVSLPPSFDCG 664
Bt 628 APSLPGINILRLHTSMYFQCWAVMCCNVPEARVFKASRSNNFYLGMLLLILFLSTMPVLYMIVSLPPSFDCG 699
Rn 493 APSLPGINILRLHTSMYFQCWAVMCCNVPEARVFKASRSNNFYLGMLLLILFLSTMPVLYMIVSLPPSFDCG 564
Clf 493 APSLPGINILRLHTSMYFQCWAVMCCNVPEARVFKASRSNNFYLGMLLLILFLSTMPVLYMIVSLPPSFDCG 564
Fc 711 APSLPGINILRLHTSMYFQCWAVMCCNVPEARVFKASRSNNFYLGMLLLILFLSTMPVLYMIVSLPPSFDCG 782
Gg 697 APCLPAINVFRLHTSMYLQCWAVMCCNVPQERVFKASRSNNFYMAMLLFILFLSTLPAVYTIVSIPPSFDCG 768
Dr 766 APCLPALNLLRLHVSMYLQCWAVMCCNVPQERVFKASGSNNFYMAMLLVILFLSTLPAIYTIVSIPPSFDCG 837
Am 645 APCLPAINIFRLHTSMYLQCWAVMCCNVPQERVFKASRSNNFYMAMLLFILFLSTLPVVYTVVSIPPSFDCG 716
Cm 699 APCLPAINVFRLHTSMYLQCWAVMCCNVPHERVFKASRSNNFYMAMLLFILFLSTLPAVYTIVSIPPSFDCG 770

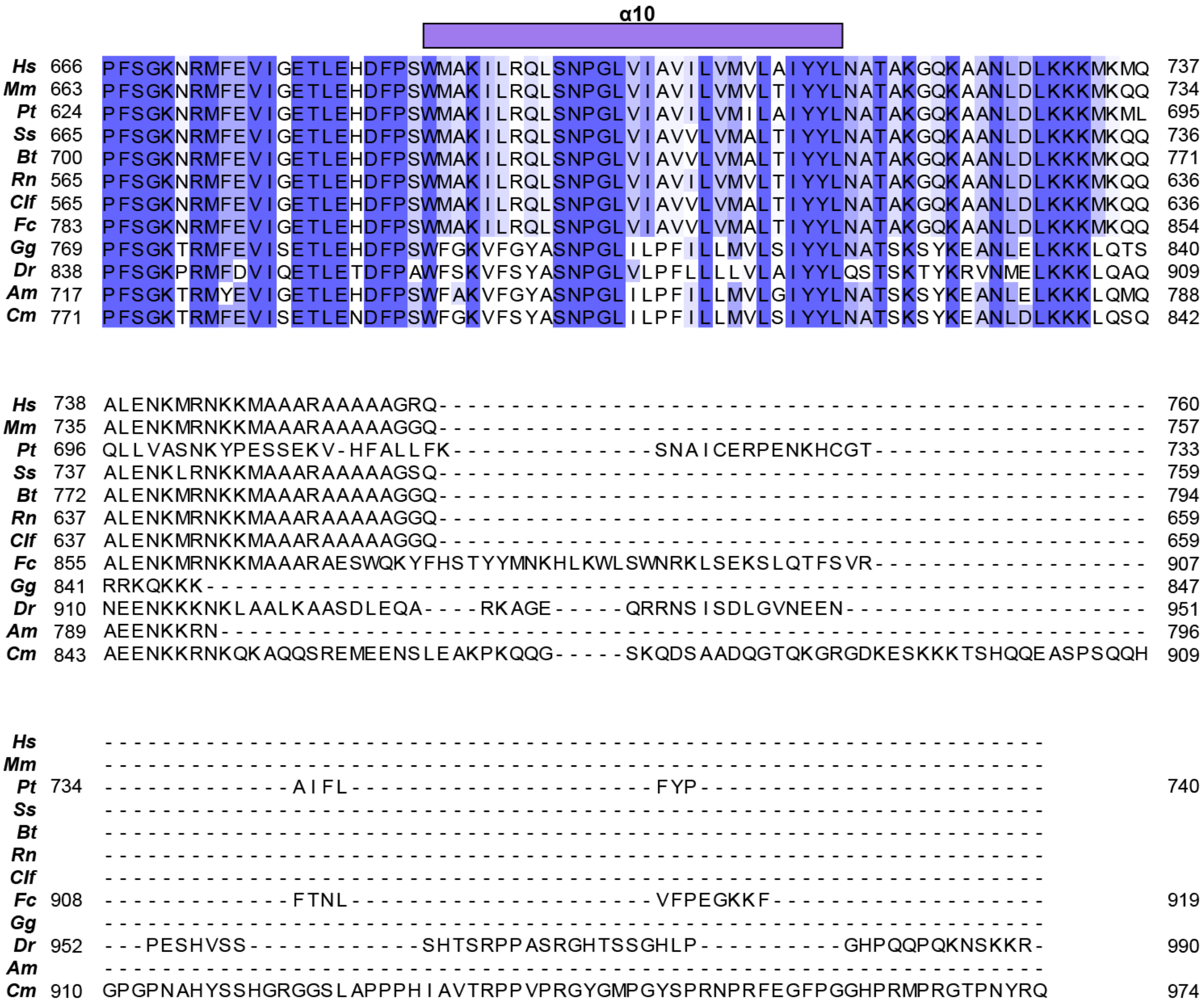


Multiple sequence alignment comparing sequences of TMC1 from 12 different species. Each sequence is colored by percent identity, where white corresponds to lowest percent identity and dark blue corresponds to highest percent identity. Red hexagons denote selected sites involved in deafness mutations, green circles denote the pore lining residues shown in Fig. S4d, and orange stars indicate hydrophobic residues that interact with lipid tails shown inf Fig S4a. Species abbreviated are as follows: *Homo sapiens* (*Hs*, NCBI Accession ID: NP_619636.2), *Mus musculus* (*Mm,* NP_083229.1), *Pan troglodytes* (*Pt*, XP_016816447.1), *Sus scrofa* (*Ss*, XP_020920511.1), *Bos taurus* (*Bt*, XP_024851786.1), *Rattus norvegicus* (*Rn*, NP_001101991.1), *Canis lupus familiaris* (*Clf*, XP_038308214.1), *Felis catus* (*Fc*, XP_023099031.1), *Gallus gallus* (*Gg*, NP_001006580.1), *Danio rerio* (*Dr*, NP_001299610.1), *Alligator mississippiensis* (*Am,* XP_014464105.1), *Chelonia mydas* (*Cm*, XP_037758993.1).
